# The Balanced Mind and its Intrinsic Neural Timescales in Advanced Meditators

**DOI:** 10.1101/2024.08.29.609126

**Authors:** Saketh Malipeddi, Arun Sasidharan, Rahul Venugopal, Bianca Ventura, Clemens Christian Bauer, Ravindra P.N., Seema Mehrotra, John P John, Bindu M Kutty, Georg Northoff

## Abstract

A balanced mind, or equanimity, cultivated through meditation and other spiritual practices, is considered one of the highest mental states. Its core features include deidentification and non-duality. Despite its significance, its neural correlates remain unknown. To address this, we acquired 128-channel EEG data (n = 103) from advanced and novice meditators (from the Isha Yoga tradition) and controls during an internal attention (breath-watching) and an external attention task (visual-oddball paradigm). We calculated the auto-correlation window (ACW), a measure of brain’s intrinsic neural timescales (INTs) and assessed equanimity through self-report questionnaires. Advanced meditators showed higher levels of equanimity and shorter duration of INTs (shorter ACW) during breath-watching, indicating deidentification with mental contents. Furthermore, they demonstrated no significant differences in INTs between tasks, indicating non-dual awareness. Finally, shorter duration of INTs correlated with the participants’ subjective perceptions of equanimity. In conclusion, we show that the shorter duration of brain’s INT may serve as a neural marker of equanimity.

## Introduction

“*The faculty of voluntarily bringing back a wandering attention, over and over again, is the very root of judgment, character, and will. … An education which should improve this faculty would be the education par excellence*.” – William James^1^.

Equanimity, observed across different cultures and spiritual traditions, is defined as “*an even-minded mental state or dispositional tendency toward all experiences or objects, regardless of their affective valence (pleasant, unpleasant or neutral) or source.”*^2^ The world’s major religions and ancient philosophies regard equanimity as one of the highest states of mind^3–7^. Distinct from mindfulness, equanimity is considered the most significant predictor of enduring happiness and well-being^2,4^. Meditation encompasses several practices from different cultural traditions, all fundamentally designed to provide insights into the true nature of the self, cultivate equanimity, and transcend suffering^3,8–15^. Therefore, equanimity is a key component of our very human basic existential states, potentially reflecting on our brain and its organization.

At the psychological level, equanimity refers to the distancing or deidentifying from internal stimuli, such as one’s own thoughts^2,16,17^. Specifically, equanimity is characterized by reduced or shorter duration of internal attention with shorter automatic reactivity to stimuli. This allows individuals to distance themselves from their own thoughts, resulting in what is described as deidentification (a term similar to cognitive dereification in the literature)^2,16–19^. In other words, stimuli and thoughts persist for a shorter duration in the mind after they are perceived, enhancing present-moment awareness, and reducing narrative and self-referential processing^2,16,17^. Overall, the scientific literature indicates that the “*primary signature of equanimity is in the temporal domain, in the form of a more rapid disengagement from… and faster return to baseline…*”^2^. In short, deidentification can be characterized by a reduced duration of internal attention to mental contents.

Another core feature of equanimity is the experience of non-dual awareness^20^, which is defined as “*a background awareness that precedes conceptualization and intention and that can contextualize various perceptual, affective, or cognitive contents without fragmenting the field of experience into habitual dualities*”^21^. A non-dual state is characterized by attenuated or dissolved boundaries between the self and others^20–31^, where one perceives reality without the usual divisions such as subject versus object or self versus other. This results in the balance and ultimately the non-distinction of internal and external attention^18,20,21,23,24,32,33^. However, the exact phenomenological and neural mechanisms underlying the state of equanimity, including both deidentification and non-dual awareness, remain yet unclear. Addressing this gap in our knowledge is the primary goal of our study.

Though we have a wide variety of subjective questionnaires to measure equanimity^16,34–39^, there is a need for more objective assessments of equanimity^2^. Electroencephalography (EEG), with its high temporal resolution, provides valuable information on ongoing brain activity, making it an important tool for investigating equanimity. Notably, studies have shown distinct EEG changes, including increases in theta and gamma power, particularly in advanced meditators^40–44^. Despite these efforts, there are currently no established neural measures to capture equanimity, including its core features like deidentification and non-duality. Consequently, the neural correlates of equanimity including its temporal features remain yet unknown. Most importantly, there is a critical need to integrate neural measures with the phenomenological features of experience and its subjective assessments of equanimity.

Since the primary signature of equanimity lies in the temporal dimension with reduced duration at the psychological level, examining duration on the neural level, as in the brain’s neural timescales, would be a promising approach to investigating equanimity. Recent evidence shows that the brain, in both humans and non-human primates, has a unimodal-transmodal hierarchy of intrinsic neural timescales (INTs)^45–52^. INTs allow for the processing of stimuli characterized by different temporal signatures, i.e., different sets of timescales. Further, research shows that these INTs provide temporal receptive windows (analogous to spatial receptive fields in the visual cortex^53–56^) that segregate and integrate inputs of varying durations^48,57–64^. Temporal segregation (separating different stimuli) is predominantly carried out by the unimodal brain regions like primary sensory cortices and has shorter INTs^46,47,62,65,66^. On the other hand, temporal integration (combining different stimuli within one neural activity) dominates in the transmodal brain regions like the lateral and medial prefrontal cortex and the default mode network (DMN) with their longer INTs^45–47,50,59,61–64,66,67^. These INTs are measured by the auto-correlation function (ACF), which is defined as a function “*that correlates a signal with copies of itself that are temporally shifted with a series of lags*^46,47,50,61^.” It is typical in research to report the autocorrelation window (ACW), defined as “*the length of time at the moment when the ACF decays to 50% of its maximum value (ACW-50)*^61,68–70^.” Measured in this way, INTs, observed during task and resting states, have been shown to play a role in perception, behavior, cognition, consciousness, and a sense of self^45–47,50,68,69,71,72^. This leaves open their role in meditation and, specifically, in mediating deidentification and non-duality as core features of equanimity.

The overarching goal of our research is to study equanimity, including both its core features, deidentification and non-duality, in terms of neural duration as the psychological observations suggest shorter duration of internal attention to mental contents. For that purpose, we study how ACW in advanced meditators differs from that in controls during different tasks and explore how these neural correlates relate to the participants’ equanimity at the psychological level. To this end, we conducted high-density (128-channel) EEG recordings during breath-watching as an internal attention task and a visual oddball paradigm^73^ as an external attention task (Figure 1a). Subsequently, we categorized these tasks along an internal-external attention gradient (Figure 1b). Furthermore, we administered the meditation depth questionnaire^74^ after the breath-watching task to evaluate participants’ state equanimity, including its two core features, namely deidentification and non-duality. Finally, we provided additional self-report questionnaires to assess non-attachment^37^ and affect balance^75^, measuring other indicators of equanimity in order to relate them to the ACW as measure of neural duration.

**Figure 1.**
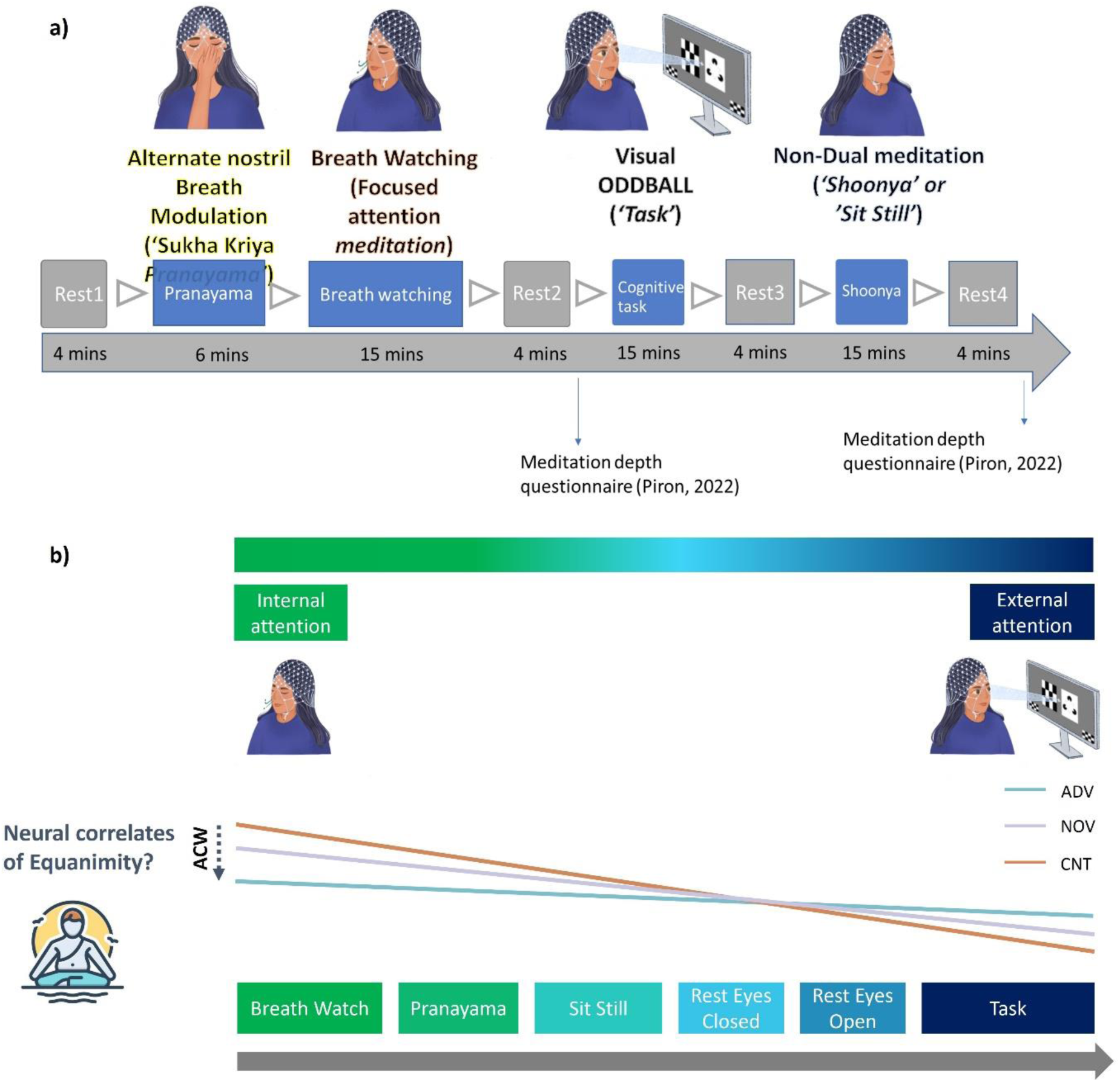
**a)** The EEG procedure consisted of four distinct sessions: pranayama, breath watching, cognitive task, and shoonya meditation. Each session included a 4-minute rest period at the beginning and end, alternating between one-minute eyes-closed and eyes-open conditions. Participants were instructed to sit still and relax during these periods. Both meditators and meditation-naïve controls practiced alternative nostril-breathing pranayama and breath watching. Controls received instructions before the study. Meditation depth during breath watching and shoonya was assessed after Rest2 and Rest4 respectively. During shoonya meditation, meditators followed the Isha tradition, while controls were instructed to sit still and given similar instructions as shoonya. **b) Study hypothesis.** Tasks were categorized along an internal-external attention gradient. We hypothesized that advanced meditators would exhibit a shorter ACW during breath watching, indicating increased temporal segregation, and a lower ACW variance indicating reduced automatic reactivity, which aligns with the fundamental feature of equanimity—disidentification from mental contents. Conversely, during external tasks, we anticipated longer ACW in advanced meditators, as they are likely to maintain internal processing, such as focusing on the breath, even during external attention-demanding activities, promoting temporal integration. Finally, we expected advanced meditators to be in a state of non-dual awareness and demonstrate a higher balance in the timescales duration across states, manifested as no significant difference or a flatter gradient in the ACW between breath watching and external task.

Our first specific aim was to investigate equanimity at the psychological level among study participants and evaluate core features of equanimity, deidentification and non-duality as well as meditation depth. Based on previous literature, we hypothesized that advanced meditators would exhibit higher levels of both state and trait equanimity, including its two core features^2,17,76^. Specifically, we hypothesized that non-duality would mediate the impact of proficiency in meditation practice on equanimity including its degree of deidentification^20^.

Our second specific aim was to characterize equanimity in terms of duration at the neural level by studying the ACW in advanced meditators and controls during both internal and external attention tasks, that is, breath-watching and visual oddball tasks, respectively. Given the nature of the internal attention task—where breath-watching involves focusing on a single stimulus (the breath) amidst competing stimuli such as mind-wandering and external distractions^32,77^—we hypothesized that advanced meditators would exhibit shorter duration in their timescales, reflected in shorter ACW (Figure 1b). This indicates greater temporal segregation (i.e., shorter ACW) (separating the breath from other stimuli) and reduced automatic reactivity (i.e., lower ACW variance), promoting a reduction in internal attention to thoughts and greater deidentification from one’s mental contents, fundamental features of equanimity. Additionally, we hypothesized that advanced meditators would be in a state of non-dual awareness and thus show a higher balance in timescales’ duration across tasks (another important signature of equanimity). This would be reflected by no significant difference, or a flatter gradient, in ACW between the internal attention demanding breath-watching and the external attention task (Figure 1b).

Our third specific aim was to address whether these neural measures of equanimity correlate with subjective measures of equanimity. We hypothesized a negative correlation between ACW and equanimity because of the shorter duration and stability of timescales at which equanimity operates on the psychological level.

We recruited advanced (5507 mean hours) and novice meditators (1637 mean hours) from the Isha Yoga tradition in India (see Table 1, Supplement for classification criteria). Isha Yoga, an international school of Yoga, is a holistic system that offers tools to promote overall well-being. Isha Yoga practices include various asanas (body postures), kriyas (breath-based techniques), and meditation (such as breath watching, Shoonya, and Samyama), among others^3,44,78,79^. Previous research on these practices has demonstrated several beneficial effects, including reduced stress and anxiety, increased mindfulness, improved immune system profiles, higher vagal tone, and enhanced well-being^78–80,80–83,83–89^. Additionally, EEG and fMRI studies have shown altered brain oscillatory dynamics and resting-state connectivity in practices promoted by the Isha tradition^44,90^. An important feature for this tradition is the cultivation of equanimity, that is to remain calm and stable, irrespective of the affective valence of situations or objects^2^. Therefore, we test this by assessing ACW in meditators and controls during internal attention (e.g., breath watching) and external attention (e.g., visual oddball) tasks as shown in figures 1 (a-b).

## Results

### Socio-demographic characteristics and meditation proficiency

The socio-demographic details of the study participants are provided in Table 1. The sample was matched for age, education, marital status, socio-economic status, and occupation.

| **Characteristics** | **Advanced**<br>**(ADV)** | **Novice**<br>**(NOV)** | **Controls**<br>**(CNT)** | **p-value** |
| --- | --- | --- | --- | --- |
| **Total sample (N)** | 42 | 33 | 28 | 0.23 |
| **Gender (M/F)** | 24/18 | 19/14 | 12/16 | 0.10 (M)<br>0.86 (F) |
| **Mean age (SD)** | 35.57<br>(6.81) | 31.66 (7.64) | 31.14<br>(6.38) | 0.01 (no<br>significant<br>differences<br>between<br>groups) |
| **Education** |  |  |  |  |
| Intermediate | 1 | 2 | 0 | 0.37 |
| Graduate | 41 | 31 | 28 | 0.10 |
| **Marital Status** |  |  |  |  |
| Married | 24 | 15 | 11 | 0.07 |
| Unmarried | 18 | 18 | 17 | 0.98 |
| **Occupation** |  |  |  |  |
| Profession | 42 | 33 | 28 |  |
| **Monthly income (Rs)** |  |  |  |  |
| 20,715-41,429 | 0 | 1 | 0 |  |
| more than 41,430 | 42 | 32 | 28 | 0.22 |
| **Meditation proficiency as mean lifetime hours (SD)** | 5507.80<br>hours<br>(2897.44) | 1637.24<br>hours<br>(1126.61) | 0 | < 0.05 |

### Psychological features of equanimity

#### Meditation depth and non-duality

We investigated the differences in the key phenomenological features of meditation among our three groups. As illustrated in Figure 2(a), the results show a significant difference in meditation depth (χ^2^_Kruskal-Wallis_(2) = 25.42, *p* = 3.02e^-06^, ɛ^2^_ordinal_ = 0.37, CI_95%_[0.25, 1.00]) and non-duality after breath-watching (χ^2^_Kruskal-Wallis_(2) = 22.94, *p* = 1.04e^-05^, ɛ^2^_ordinal_ = 0.33, CI_95%_[0.21, 1.00]) between the three groups. Advanced meditators showed the highest levels in both meditation depth and non-duality, followed by novices, with controls reporting the lowest levels. The effect sizes were large.

**Figure 2.**
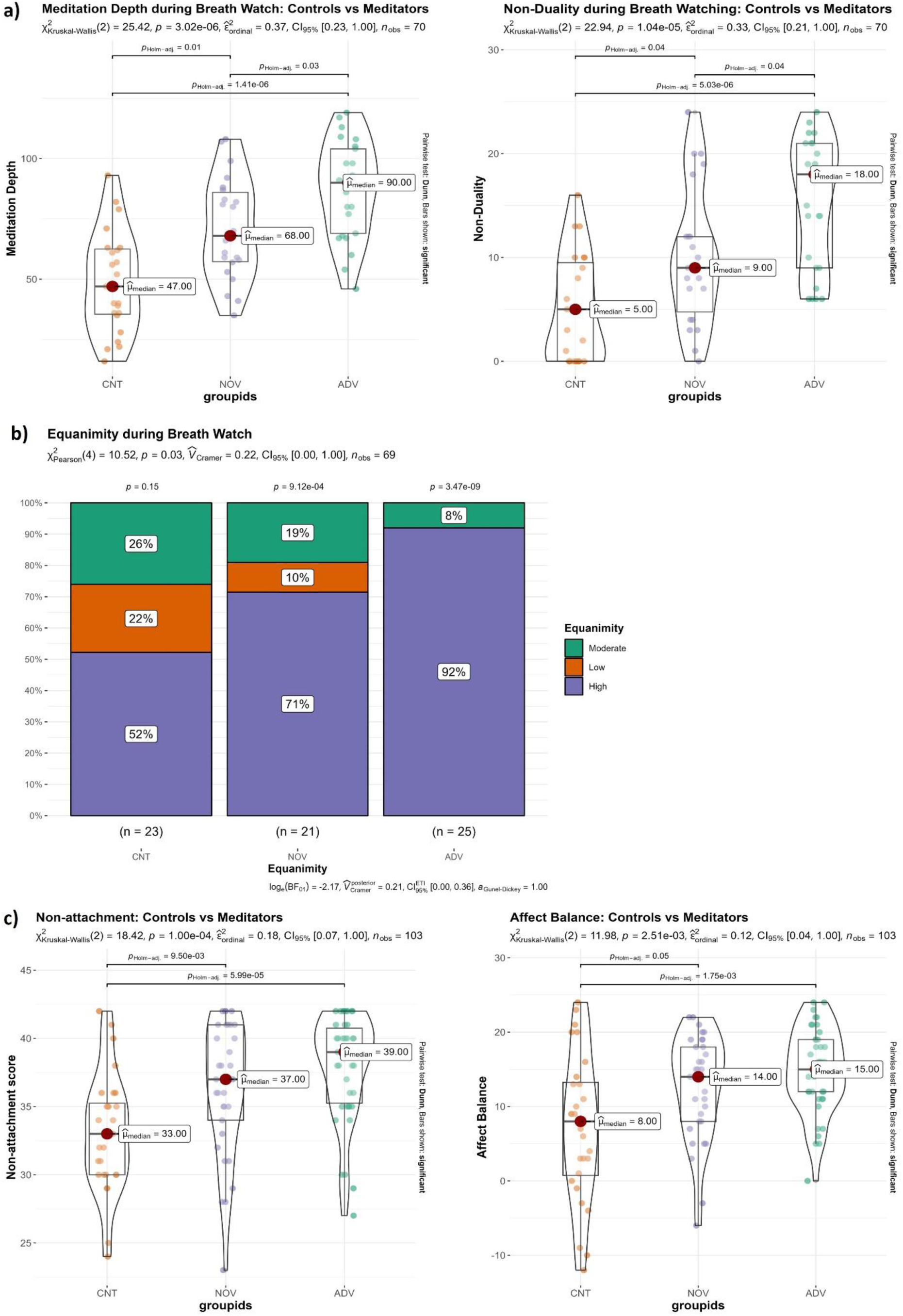
**a)** Meditation depth (left) and non-duality (right) during breath watching **b)** Equanimity during breath watch and **c)** Non-attachment (left) and affect balance (right). CNT – Controls, NOV – Novice meditators, ADV – Advanced meditators

#### Deidentification and Equanimity

Item 2 in our meditation depth scale directly refers to equanimity, which we therefore compared among the groups. As shown in Figure 2(b), the results show a significant difference between the groups in state equanimity during breath watch (χ^2^_Pearson_(4) = 10.52, *p* = 0.03, V_Cramer_ = 0.22, CI_95%_[0.00, 1.00]). High equanimity was reported by 92% of advanced meditators, with none reporting low equanimity. In contrast, 52% of controls reported high equanimity, while 22% reported low equanimity. Additional support for greater equanimity among advanced meditators is found when we understand equanimity in a wider sense by including different items from our meditation scale, like 4-item-equanimity, with item 2 + 3 other items from the meditation depth scale and 8-item-equanimity, with item 2 + 7 items from the meditation depth scale, which also revealed a significant difference between the groups (see Supplement Figure 1 and 2).

#### Non-attachment and Affect Balance

In addition to measuring the actual state directly after breath watching with our meditation scale, we also investigated subjects’ trait features with other scales. For this, we assessed non-attachment and affect balance between the groups. Results (figure 2c) show a significant difference between the groups in both non-attachment (χ^2^_Kruskal-Wallis_(2) = 18.42, *p* = 1.0e^-04^, ɛ^2^_ordinal_ = 0.18, CI_95%_[0.10, 1.00]) and affect balance (χ^2^_Kruskal-Wallis_(2) = 11.98, *p* = 2.51e^-03^, ɛ^2^_ordinal_ = 0.12, CI_95%_[0.05, 1.00]). The effect sizes were large for non-attachment and medium for affect balance. Advanced meditators reported highest scores for both non-attachment and affect balance. Further, we observed higher positive and lower negative feelings, reflecting questions of the affect balance scale, in advanced meditators (Supplementary Figure 3).

#### Relationship between the different psychological measures

Based on our results that advanced meditators show higher levels of both state and trait equanimity, we next investigated the relationship among its different features. To do so, we conducted a mediation analysis. As depicted in Figure 3, the findings revealed that non-duality serves as a mediator in the relationship between lifetime hours of meditation and equanimity (figure 3, top). The direct path from meditation lifetime hours to equanimity (c’ in the figure) was non-significant (z = 1.659, *p* = 0.097, CI_95%_[0.000, 0.000]). While the paths “a” (z = 3.469, *p* = 0.001, CI_95%_[0.001, 0.002]) and “b” (z = 5.254, *p* = 0.000, CI_95%_[0.052, 0.113]) were significant. The path “a*b” was also significant (z = 2.653, *p* = 0.008, CI_95%_[0.000, 0.000]). This suggests a full mediation effect.

**Figure 3:**
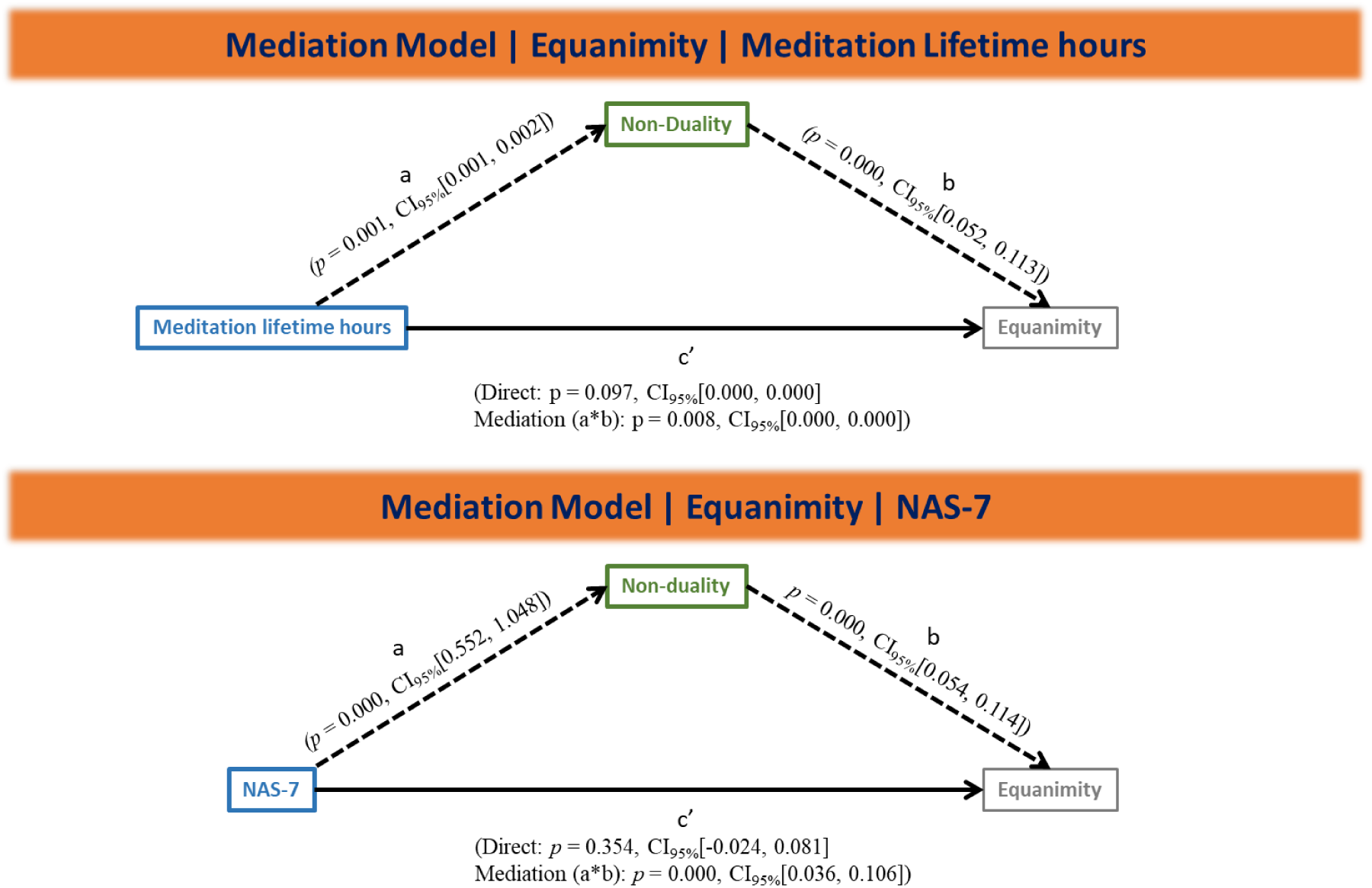
Mediation analysis showed that non-duality (mediator) mediates the effect of meditation lifetime hours (top) and non-attachment (bottom) on Equanimity.

Similarly, results showed that non-duality mediates the effect of non-attachment on equanimity (figure 3, bottom). The direct path from non-attachment to equanimity (c’ in the figure) was non-significant (z = 0.926, *p* = 0.354, CI_95%_[-0.024, 0.081]). While the paths “a” (z = 6.237, *p* = 0.000, CI_95%_[0.552, 1.048]) and “b” (z = 5.363, *p* = 0.000, CI_95%_[0.054, 0.114]) were significant. The path “a*b” was also significant (z = 3.685, *p* = 0.000, CI_95%_[0.036, 0.106]), indicating a full mediation effect. To validate the models, we exchanged the factors, but no found no evidence of significant differences.

To further investigate the association between non-duality and equanimity, we merged the data from both meditators and controls, performed a median-split of the data, and categorized participants into four groups: Low Non-duality Low Equanimity, Low Non-duality High Equanimity, High Non-duality Low Equanimity, and High Non-duality High Equanimity (see Supplementary Figure 4). Significant differences were observed among these groups, with most participants (44%) falling into the High Non-duality High Equanimity category, and the fewest (4%) in the High Non-duality Low Equanimity category. Additionally, 29% of participants were classified into the Low Non-duality High Equanimity group. Significant differences were also found between these four groups in terms of meditation lifetime hours (see Supplementary Figure 5) and non-attachment (see Supplementary Figure 6).

Together, our data show high levels of equanimity and its various features, such as deidentification, non-attachment, and affective balance, in the advanced meditators. Moreover, we demonstrate that the effects of meditation lifetime hours and non-attachment on equanimity, including deidentification, are fully mediated by non-duality.

### Duration as neural correlate of equanimity

#### Autocorrelation window (ACW) during Breath Watch: Controls vs Advanced meditators

Next, we investigated internally and externally oriented durations of neural processing and balance in timescales duration across tasks as key features of equanimity at the neuronal level. Duration can be operationalized by the autocorrelation window (ACW) during a task requiring high internal attention, such as breath watching. We were thus interested in investigating if ACW mean (and ACW stability as measured by the variance calculated across its sliding windows) values during breath watch would differ between controls and advanced meditators. To examine this, we carried out a Mann-Whitney test between these two groups. We found a statistically significant shorter ACW mean (*W*_Mann-Whitney_ = 306.00, *p* = 0.01, r_rank-biserial_ = -0.38, CI_95%_[-0.60, -0.11]) (Figure 4a) and a lower variance (*W*_Mann-Whitney_ = 294.50, *p* = 0.03, r_rank-biserial_ = -0.33, CI_95%_[-0.57, -0.05])) (Supplementary Figure 7) in advanced meditators compared to controls. The effect sizes were large. We also investigated this during external task, but we did not find any ACW difference between the groups (Supplementary Figure 8). In sum, the duration of neural activity, e.g., ACW during high internal attention as in breath watch, is shorter in advanced mediators than in controls. Moreover, we observed more stability of the ACW, as measured by its variance with a sliding window approach (see supplementary material) in the advanced meditators reflecting their higher degree of temporal stability.

**Figure 4.**
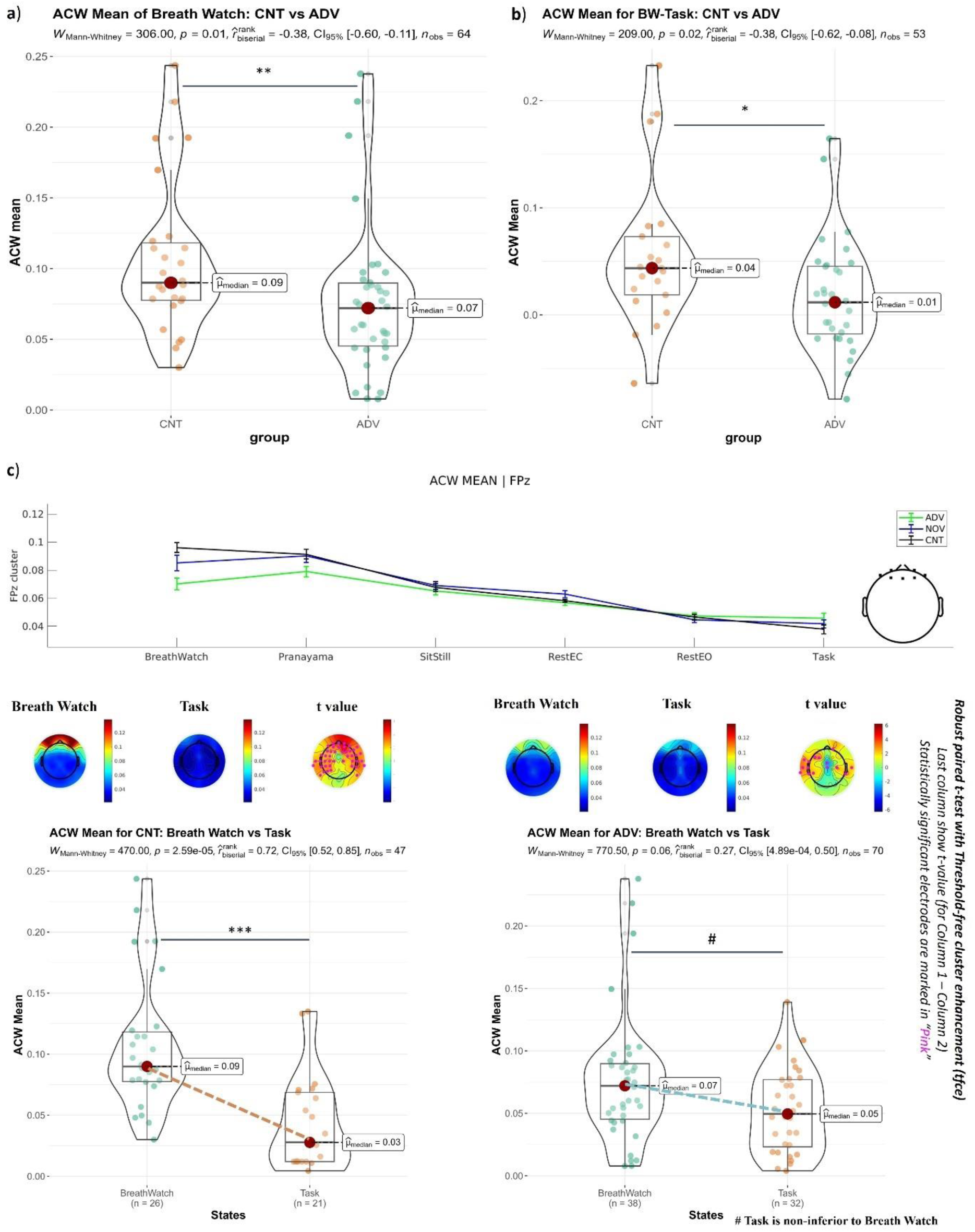
**(a):** ACW mean during breath watch between controls and advanced meditators. **b):** Plot shows comparison between controls and advanced meditators in difference scores of ACW mean between breath watch and task (BW-Task). **c)** The top panel shows the line plot for the ACW mean (FpZ cluster, top right) along the internal-external attention gradient. The middle panel shows the topoplots of ACW mean during breath watch and task for controls (left) and advanced meditators (right). The last column of the topoplots shows t-values (column 1 – column 2) and statistically significant electrodes are marked in pink. The bottom panel shows the ACW mean for controls (left) and advanced meditators (right) for breath watch vs. task. Statistical significance: * < 0.05, ** < 0.01, *** < 0.001.

#### ACW for Breath Watch vs Task in controls and advanced meditators

We next operationalize the non-duality by comparing the differences in ACW along a gradient of high internal and external attention tasks, that is, from breath watch to oddball task. We assessed how participants processed tasks along the internal-external attention gradient (Figure 1b) by calculating the ACW mean for all tasks (Figure 4c, top panel). Results indicated a flatter gradient for the ACW mean in advanced meditators compared to other groups. We then conducted a Mann-Whitney test comparing the ACW mean for breath watch and visual oddball between controls and advanced meditators. The analysis revealed a significant difference, with a very large effect size, in the ACW mean between breath watch and task for controls (*W*_Mann-Whitney_ = 470.00, *p* = 2.59e^-05^, r_rank-biserial_ = 0.72, CI_95%_[0.52, 0.85]) (Figure 4c, bottom panel, left). Conversely, we found no evidence for a difference in ACW mean between breath watch and task in advanced meditators (*W*_Mann-Whitney_ = 770.50, *p* = 0.06, r_rank-biserial_ = 0.27, CI_95%_[4.89e^-04^, 0.50]) (Figure 4c, bottom panel, right). Topo-plots of the ACW mean indicated significant differences between breath watch and task in the entire anterior regions of the brain for controls (Figure 4c, middle panel, left), whereas few electrodes showed significant differences for advanced meditators (Figure 4c, middle panel, right). Similar results were found for ACW variance (Supplementary Figure 9). Further, results showed a significant difference between controls and advanced meditators in the breath watch and task difference scores of the ACW mean (*W*_Mann-Whitney_ = 209.00, *p* = 0.02, r_rank-biserial_ = -0.38, CI_95%_[-0.62, -0.08] (Figure 4b).

For additional validation, we conducted a Mann-Whitney test comparing the ACW mean (and variance) between rest eyes closed (RestEC) and breath watch in both controls and advanced meditators. The analysis revealed a significant difference, with a large effect size, between rest eyes closed and breath watch in controls for the ACW mean (*W*_Mann-Whitney_ = 535.00, *p* = 3.16e^-03^, r_rank-biserial_ = 0.47, CI_95%_[0.20, 0.68]) (Supplementary Figure 10, left). However, no evidence of a difference between rest eyes closed and breath watch for ACW mean was found in advanced meditators (*W*_Mann-Whitney_ = 735.00, *p* = 0.90, r_rank-biserial_ = 0.02, CI_95%_[-0.24, 0.27]) (Supplementary Figure 10, right). We observed similar results for ACW variance (Supplementary Figure 11).

Together, we show that the advanced meditators exhibit less ACW differences between high internal and high external task conditions than the controls and the novices. Moreover, shortening their ACW during the internal attention state, e.g., breath watch, puts it in a similar range as the one during the external attention state, with both internal and external states no longer being significantly different from each other. Moreover, variability of ACW was also decreased in both conditions in the advanced mediators again reflecting their increased temporal stability.

### From neural duration to equanimity

#### Correlation between ACW mean (and variance) and Meditation depth (and non-duality)

Our final step consists in relating the neuronal measures of duration and non-duality, e.g., ACW during internal and external attention tasks, to their psychological measures, e.g., the self report sales. How do ACW values correlate with participants’ self-report measures? Our findings show a negative correlation between meditation depth and the ACW mean of breath watching (Ρ_Spearman_ = -0.24, *p* = 0.0473) (Figure 5a). Additionally, meditation depth and the ACW variance were negative correlated (Ρ_Spearman_ = -0.3, *p* = 0.0149) (Figure 5b). Similarly, we observed a negative correlation between non-duality and the ACW mean (Ρ_Spearman_ = - 0.28, *p* = 0.0211) (Figure 5c) (as well as a negative correlation between non-duality and the ACW variance (Ρ_Spearman_ = -0.3, *p* = 0.0177)) (Figure 5d). Further validation came from negative correlations we observed between meditation lifetime hours and ACW mean (and variance) (Supplementary Figure 15). Together, these findings show that shorter ACW mean, and its lower variability are related to increased meditation depth and non-duality.

**Figure 5:**
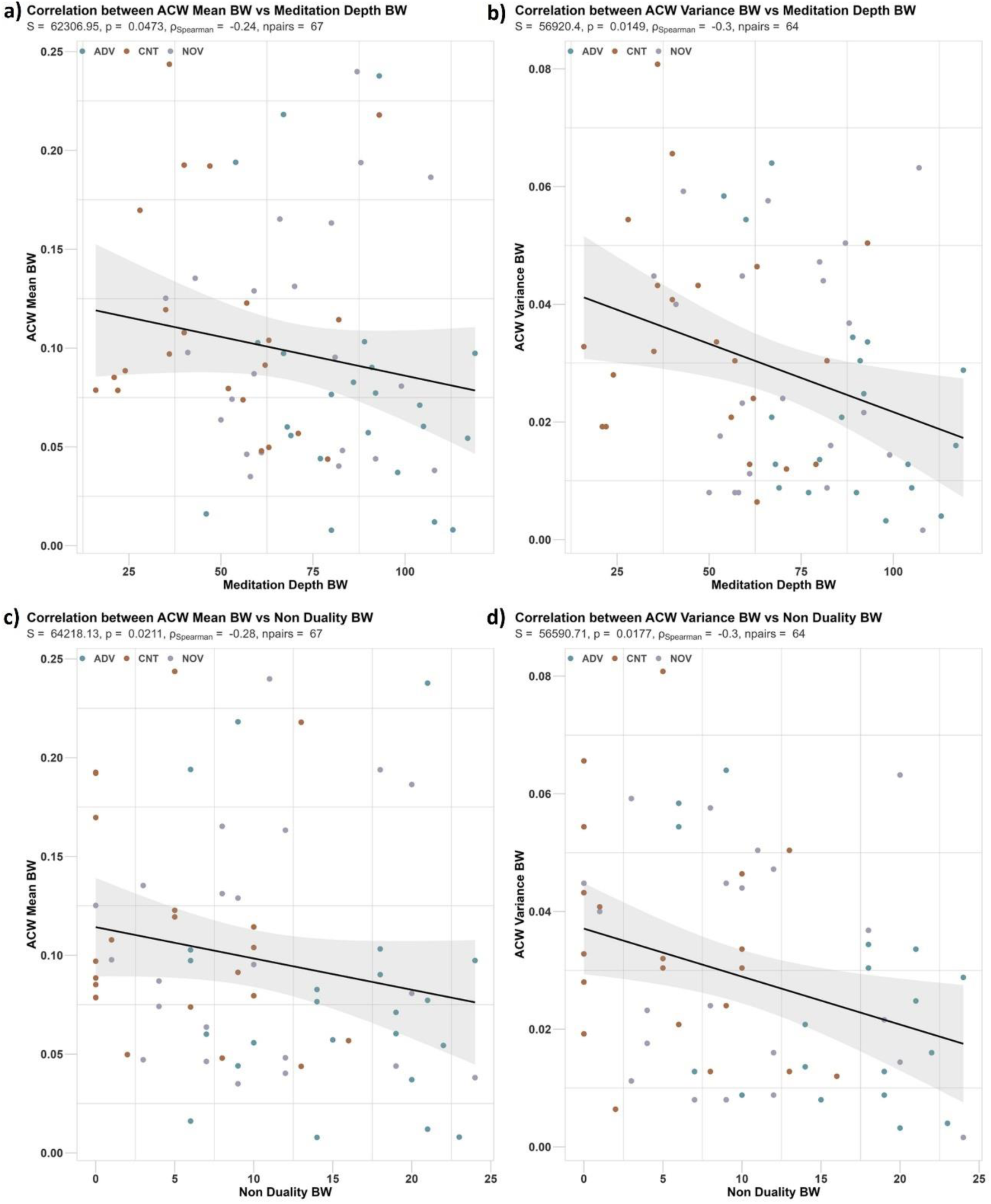
**(a)** Correlations between ACW mean and breath watch, **b)** ACW variance and breath watch, **c)** ACW mean and non-duality, and **d)** ACW variance and non-duality. Shadows show 95% CI intervals.

#### Correlation between ACW Mean BW-Task and Meditation depth (and non-duality)

We then investigated the relationship between the difference scores of ACW mean for breath watch and task (BW-Task) with meditation depth and non-duality. Our results showed a negative correlation between meditation depth and ACW mean BW-Task (Ρ_Spearman_ = -0.29, *p* = 0.0331) (Figure 6a). Similarly, a negative correlation was observed between non-duality and ACW mean BW-Task (Ρ_Spearman_ = -0.31, *p* = 0.0227) (Figure 6b). These findings suggest that the neural measure of non-duality, e.g., internal-external ACW duration difference, relates to both meditation depth and non-duality on the psychological level.

**Figure 6.**
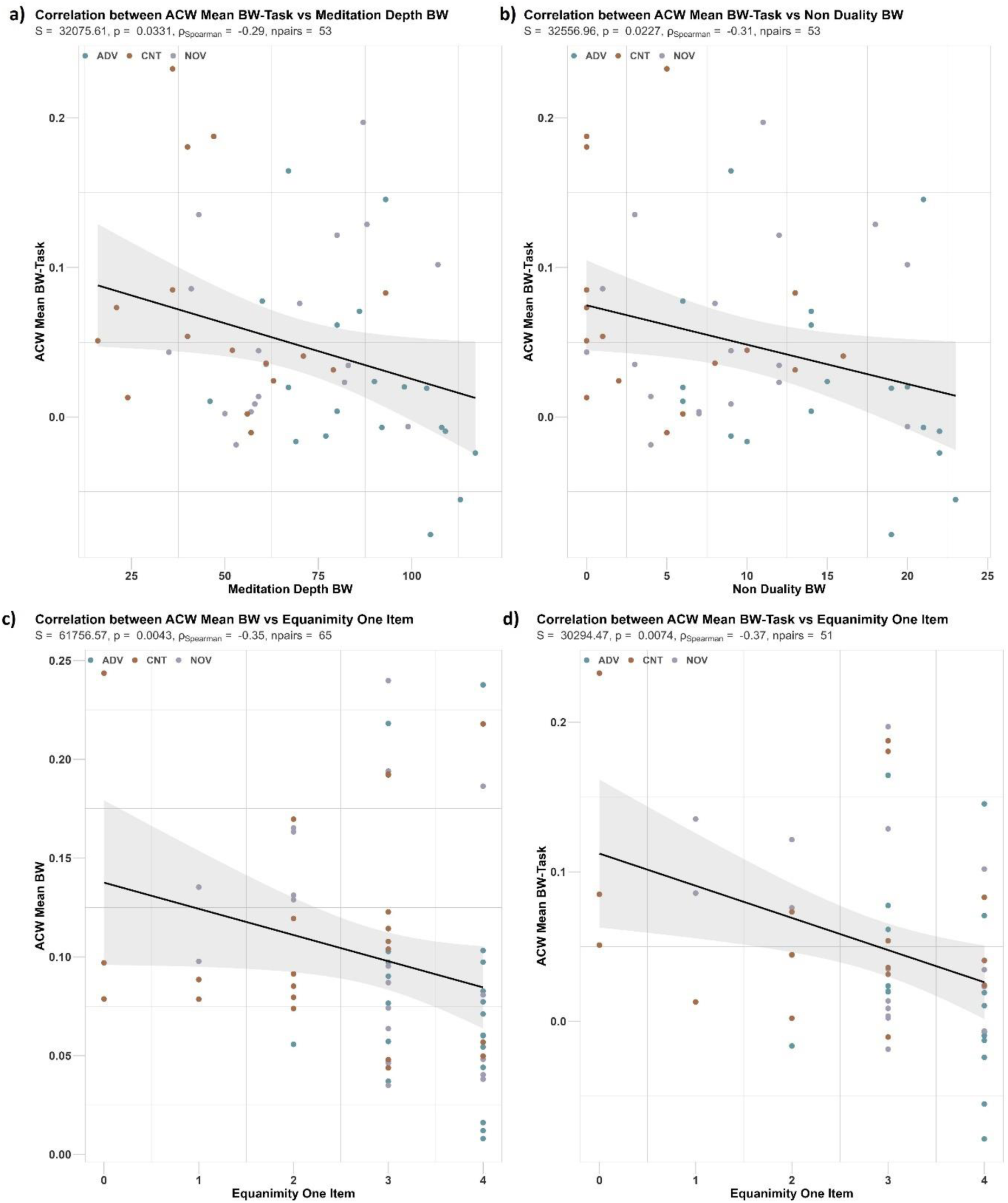
**(a):** Correlations between ACW mean BW-Task and Meditation depth, **b)** ACW mean BW-Task and non-duality, **c)** ACW mean BW and Equanimity One item, and **d)** ACW mean BW-Task and Equanimity One item. Shadows show 95% CI intervals.

#### Correlations between ACW and Equanimity

Next, we asked whether the ACW on the neural level is related directly to equanimity on the psychological level. Our results show negative correlations between equanimity and ACW mean of breath watching (Ρ_Spearman_ = -0.35, *p* = 0.0043) (Figure 6c), ACW variance of breath watching (Ρ_Spearman_ = -0.38, *p* = 0.0019 (Supplement Figure 12, left), ACW mean of BW-Task (Ρ_Spearman_ = -0.37, *p* = 0.0074) (Figure 6d), and ACW variance of BW-Task (Ρ_Spearman_ = -0.43, *p* = 0.0136) (Supplement Figure 12, right). As an additional validation, we carried out correlations between 4-item and 8-item-equanimity and ACW. We observed a negative correlation for all measures (Supplementary Figures 13 and 14). Together, these findings suggest a relation of ACW, during both internal attention and internal-external attention differences, with equanimity, as supported by various levels of analyses and scales. A shorter ACW mean and lower ACW variability are related to higher levels of equanimity. This supports our assumption that shorter neural duration and its higher temporal stability are key for equanimity.

#### Mediation models

We finally asked for the relationship among the different neural and psychological measures, that is, whether the ACW on the neuronal level affects equanimity through the mediation of non-duality, given that the latter is key in mediating the former on the psychological level (see above). As depicted in Figure 7, our findings showed that non-duality serves as a mediator in the relationship between ACW mean BW-Task (difference score) and equanimity (figure 7, top). The direct path from ACW mean BW-Task to equanimity (c’ in the figure) was non-significant (z = -1.451, p = 0.147, CI_95%_[-7.954, 0.990]). However, the paths “a” (z = - 2.166, p = 0.030, CI_95%_[-60.648, -2.820]) and “b” (z = 4.807, p = 0.000, CI_95%_[0.053, 0.125]) were significant. The path “a*b” was also significant (z = -1.975, p = 0.048, CI_95%_[-5.892, - 0.271]). This suggests a full mediation effect. Similarly, our results showed that non-duality mediates the relationship between ACW variance of breath watch and equanimity (figure 7, bottom). To validate the models, we exchanged the factors, but no found no evidence of significant differences. Together, we can now also see the key role of non-duality in mediating the relation between ACW and equanimity on the combined neural and psychological level. This further supports our assumption that a shorter and more internal-external balanced neural duration plays a key role in the neural connection to equanimity through non-duality.

**Figure 7:**
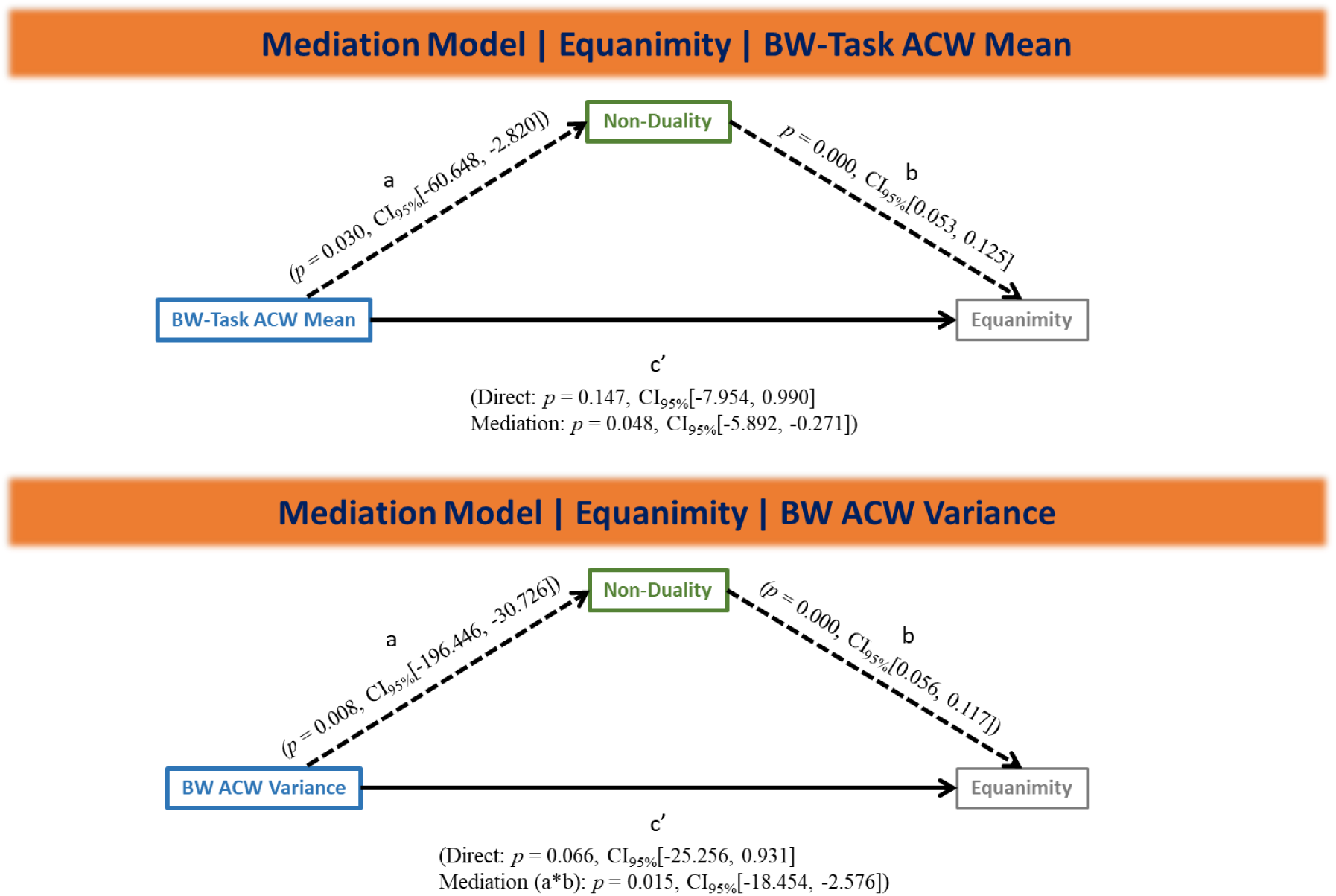
Mediation analysis showed that non-duality (mediator) mediates the effect of BW-Task ACW Mean (top) and BW ACW variance (bottom) on Equanimity. BW: Breath Watching.

## Discussion

We investigated the psychological and neural features of equanimity associated with Isha meditation practice. Advanced meditators exhibited greater state and trait equanimity compared to other groups, consistent with existing literature^2,17,76^. We have studied the brain’s intrinsic neural timescales^46,47^, measured using the auto-correlation window (ACW), to objectively assess equanimity. We found shorter and more stable ACW duration during an internal attention breath-watching task in advanced meditators, suggesting shorter and consistent processing duration, thereby facilitating de-identification with internal mental contents. Additionally, advanced meditators showed no significant difference in ACW between the breath-watching task and the visual oddball task, indicating non-duality. Finally, we found a negative correlation between ACW and equanimity, with non-duality mediating these effects. Overall, our research demonstrates that duration and its stability on the neural level, as measured by the ACW mean and variance, characterize de-identification and non-duality, respectively, as core features of equanimity in advanced meditators. This suggests that shorter and more stable duration on both neural and psychological levels is a core feature of equanimity, including de-identification and non-duality (Figure 8).

**Figure 8:**
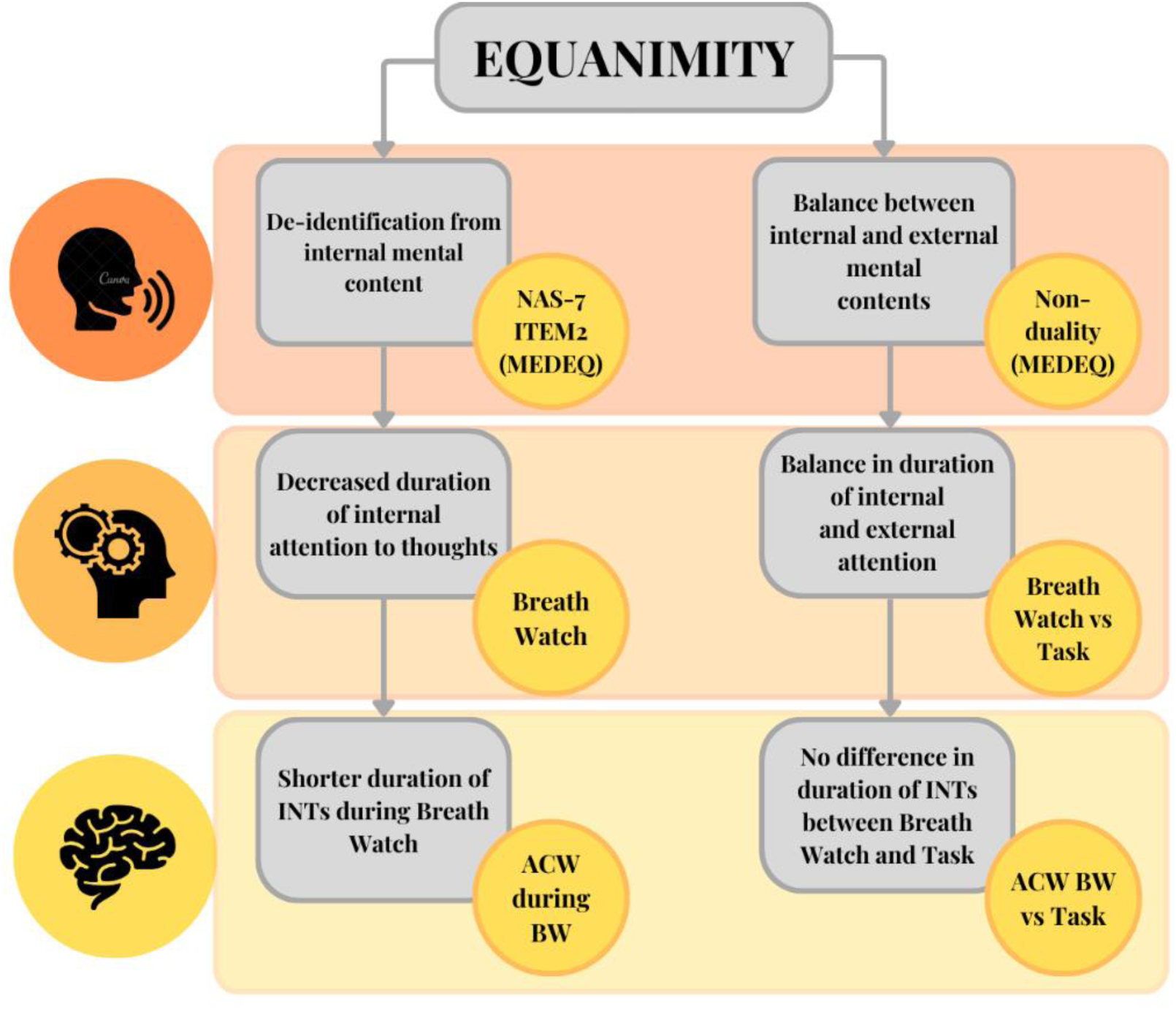
Our study links the psychological to neural indices of equanimity in advanced meditators by highlighting three distinct levels of analysis: psychological, attentional, and neural. Equanimity is defined by two main features: (1) deidentification from internal mental content and (2) balance between internal and external contents. From a psychological perspective (orange box), we assessed the reduced duration which characterizes the deidentification from internal mental content (on the left) using the NAS-7 and item 2 of the Meditation Depth Questionnaire (MEDEQ). To examine this feature on the attentional level (light orange box), we designed an experiment involving an internal attention task, specifically breath watching. On the neural level (yellow box), we found reduced duration of intrinsic neural timescales (INTs), indicated by shorter autocorrelation windows (ACW) during breath watching. Moreover, we psychologically (orange box) assessed the balance of durations assigned to internal and external contents (on the right) using the non-duality cluster of the MEDEQ. Attentional balance (light orange box) was investigated by comparing the internal attention task (breath watching) with an external attention task (visual oddball). Neurally (yellow box), we did not find any significant difference in ACW between breath watching and the visual oddball task in advanced meditators. The comprehensive approach we adopted allowed us to connect psychological assessments with neural indicators, demonstrating how equanimity manifests in experienced meditators across different levels of analysis. NAS-7: Non-attachment scale, MEDEQ: Meditation Depth Questionnaire, BW: Breath Watch, INTs: Intrinsic Neural Timescales.

We found greater equanimity during breath-watching in advanced meditators compared to other groups. In breath-watching meditation, practitioners focus attention on breath while ignoring distractions, such as mind-wandering^3,19,77,78,91^. Reduced mind wandering and increased attention focus have been consistently reported in advanced meditators from different traditions^77,90,92–100^. Increased duration of attention on breath, which leads to de-identification with internal mental contents, could promote more steady windows of attention, leading to greater equanimity^2,19,76^. An increase in state equanimity following a 30-minute breathing meditation practice compared to an active control condition^76^ has been reported. Further, an fMRI study on expert Tibetan Buddhist meditators demonstrated that they could achieve sustained attention with fewer attentional networks than novices^101^, suggesting that extensive attentional training leads to a state where attention flows effortlessly towards the object of focus, enabling a non-dual state of awareness—another core feature of equanimity^20,21,24^. Overall, studies show that meditation, through attentional training and meta-awareness, promotes deidentification with one’s thoughts and non-duality^18,19,102–104^. Furthermore, advanced meditators demonstrated greater depth of meditation and non-duality, as measured by the meditation depth questionnaire. Phenomenological studies from other labs have also reported similar experiences of non-duality during various forms of meditation^20,21,24^. Overall, our findings at the psychological level indicate that advanced meditators exhibit significantly greater equanimity, characterized by increased deidentification and non-duality.

Having identified greater equanimity at the psychological level in advanced meditators, our next question was whether this would also be reflected at the neural level. We observed shorter ACW during breath-watching in advanced meditators, suggesting they use shorter neural timescales when practicing breath-watching. Neural timescales shorten and lengthen with the segregation and integration of stimuli, respectively^46–48,50^. A focused breath-watching requires the segregation of the breath from distracting stimuli, such as thoughts, which entails reduced internal attention to mental contents, or deidentification with internal stimuli. Further, topographic plots showed shorter ACW in transmodal regions in advanced meditators, further supporting the effortless and deidentified nature of breath-watching in this group. In contrast, controls exhibited longer ACW, indicating that they process the breath alongside other unwanted stimuli like thoughts, thus prolonging the processing duration. Lower meditation depth in controls supports this.

Additionally, we found no significant differences in ACW between internally oriented breath-watching and the externally oriented task in advanced meditators, suggesting a non-dual state. During the external task, no significant differences in ACW mean were observed between advanced meditators and controls, indicating similar processing of external tasks. However, controls processed the internal task very differently. For controls, ACW lengthens during the more internal breath-watching and shortens during the external task. This results in a steeper ACW slope along the internal-external attention gradient for controls reflecting dual mode in internal and external processing. In contrast, advanced meditators exhibited similar ACW in both conditions, resulting in a flatter slope, indicating non-duality, a core feature of equanimity. The absence of significant differences between the resting state and breath-watching as well as between the resting state and the external task in advanced meditators further supports their experience of a non-dual state.

The final line of evidence supporting the key role of neural duration for deidentification and non-duality as core features of equanimity comes from the significant correlations we observed between the psychological and neural measures of equanimity. Specifically, shorter ACW durations during breath-watching and smaller ACW differences between breath-watching and task conditions were associated with greater meditation depth, non-duality, and equanimity. Moreover, non-duality mediated the effect of lifetime meditation hours on equanimity. The absence of a direct effect (c’ in Figure 3 and 7) suggests that an increase in lifetime meditation hours, without a corresponding experience of non-duality, may not lead to enhanced equanimity. Overall, our findings imply that individuals, through meditative training, can experience various events from a non-dual state of awareness, rather than habitually categorizing everything into dualities such as self versus other, good versus bad and internal versus external. This finding aligns with other studies and reports, which indicate a reduction in dualistic distinctions in advanced meditators^20–28,31,33,105,106^.

Equanimity, observed cross-culturally, is considered one of the highest states of mind and is part of the phenomenal repertoire of humanity^3,5–7,15^. The Hindu scripture Bhagavad Gita defines equanimity as “*Samatvam Yoga Uchyate,”* which translates to “*Equanimity is Yoga.”*^5^ In other words, to experience Yoga, which means union between the “self” and the “other,” one must be in a state of mental unperturbability or equanimity. Buddhism considers *Upekkha* to be one of the Four Immeasurable Minds, or four sublime qualities of the human mind. Upekkha is translated as equanimity, even-mindedness, nonidentification, nonattachment, non-discrimination, inclusiveness, or letting go^6,107^. Similarly, it is related to *Samayika* in Jainism, *Menuhat ha-Nefesh* in Judaism, *Wu Wei* in Taoism, and the virtue of *Temperance* in Christianity.

The experience of equanimity has significant implications for individual happiness and well-being^2,4,38^. While there has been considerable research on mindfulness and its various health effects, recent studies suggest the need to move beyond mindfulness and look at equanimity as a crucial predictor of enduring happiness and well-being^2,4,17,76^. In this context, our findings indicate that equanimity is also part of the neural repertoire of humanity, with meditation serving as a valuable tool for cultivating this state. By demonstrating that advanced meditators show equanimity at both psychological and neural levels, our study underscores the value of integrating meditative practices into mental health and well-being programs. Meditation, a trainable skill, helps to cultivate resilience against everyday stressors and maintain a non-reactive, balanced state of mind^10,19,20,77,102,108–112^. As the prevalence of mental health issues increase globally^113–117^, qualities such as non-duality and deidentification from thoughts that characterize equanimity offer a pathway to achieving a deeper sense of peace and stability.

## Limitations

While ACW values suggest a non-dual state in advanced meditators during external tasks, the absence of a psychological measure of duration in this state is a limitation of the current study. Further, future research should incorporate paradigms that elicit stress and emotions to determine whether the equanimous state in advanced meditators is affected by such events.

## Conclusion

Our study introduces a novel psychological and neural view of equanimity in advanced meditators. We first show that deidentification and non-duality are indeed core features of equanimity in advanced meditators, with non-duality mediating deidentification. Our neural findings show that shorter ACW duration serves as an index of deidentification with one’s thoughts and a flatter ACW slope along the internal-external attention gradient provides a neural index of non-duality. Being strongly supported by correlation and mediation findings, we suggest that the shorter neural duration and higher stability of the brain’s timescales, e.g., ACW mean and variance, in advanced meditators are key in mediating their experience of non-duality and deidentification as core features of equanimity.

## Acknowledgements

We express thanks to the volunteers of the Isha Foundation for their support in participant recruitment. We extend our gratitude to Madhumita Das, Illustrator and graphic designer, Vistar Arts, for the illustrations provided. Finally, we sincerely thank Maa Vama, Isha Foundation, for all her support and guidance in conducting this research.

## Methods

### Participants

103 healthy adults (Table 1) were recruited and categorized into three groups: advanced meditators (ADV), novice meditators (NOV), and meditation-naïve controls (CNT). The classification criteria are provided in Table 1 of the Supplementary File. Meditators were recruited from Karnataka, India, with assistance from the Isha Foundation, while control participants were recruited locally in Bengaluru, Karnataka. Advanced meditators (n = 42, 18 females) had a mean (SD) age of 35.57 (6.81) years and a mean (SD) lifetime Isha Yoga practice duration of 5507.80 (2897.44) hours. Novice meditators (n = 33, 14 females) had a mean (SD) age of 31.66 (7.64) years with a mean (SD) lifetime practice duration of 1637.24 (1126.61) hours in Isha Yoga. Meditation-naïve controls (n = 28, 16 females) had a mean (SD) age of 31.14 (6.38) years and no prior exposure to meditation or yoga schools; they were matched for age, education, and socio-economic background (Table 1). All participants were healthy or under stable non-psychoactive medication. Further details on inclusion and exclusion criteria can be found in Table 2 of the Supplementary File. This study received approval from the NIMHANS Human Research Ethics Committee (NIMH/DO/ETHICS SUB-COMMITTEE MEETING/2018), and participants provided written informed consent before participation. No monetary compensation was provided.

### Procedure

EEG studies were carried out at the Centre for Consciousness Studies, Department of Neurophysiology at NIMHANS. Participants completed self-report questionnaires (psychometrics) either before or after EEG recordings, with the order counterbalanced to reduce potential order effects. The EEG procedure consisted of four distinct sessions: alternate nostril breathing pranayama, breath watching, cognitive task, and shoonya meditation (refer to Figure 1). Both meditators and meditation-naïve controls underwent these four sessions. Each session included a 4-minute rest period at the beginning and end, alternating between one-minute eyes-closed and eyes-open conditions. Participants were instructed to sit still and relax during these periods. Controls received instructions before the study. Meditation depth during breath watching was assessed after Rest2. During shoonya meditation, meditators followed the Isha tradition, while controls were instructed to sit still and given similar instructions as shoonya. Meditation depth during shoonya was assessed after Rest4. Specific instructions provided to participants can be found in the supplementary file.

### Cognitive Task for external attention

We utilized a novel paradigm called ANGEL (Assessing Neurocognition via Gamified Experimental Logic) in our study^73^. This paradigm is based on the visual oddball paradigm, but it includes various gaming elements to increase engagement and allow for the investigation of decision-making in a variety of contexts. ANGEL involves the simultaneous presentation of multiple audio and visual stimuli, challenging participants to make decisions amidst various distractions. This paradigm allows the multiple assessment of 10 different event-related potentials (ERPs). The task consists of 448 trials (16 blocks, each comprising 25 stimulus trials and 3 baseline trials) and lasts approximately 15 minutes. Each trial begins with a pre-stimulus phase featuring a white plus sign used for fixation against a grey background with checkerboards displayed on both sides. There are four possible types of novel images: a Mooney face or a distorted version thereof (randomly selected from a set of 30 each), and a Kanizsa triangle or a distorted image comprising the same components as the Kanizsa triangle (randomly chosen from a set of two each). In each block, the oddball paradigm is implemented with different versions, featuring the presentation of salient image types as follows: One of the four salient image types is displayed on one side of the fixation sign for 80% of the time, constituting the frequent image category, while the other two image types are shown on the opposite side, totalling 20% of the time (10% for each type) and forming the rare category. Participants received performance feedback at the end of every two blocks to keep them motivated, and instructions and practice sessions were provided prior to the main session

### Isha Yoga

Isha Yoga offers tools for inner well-being without subscribing to any specific philosophy or belief system^3^. It encompasses practices such as Hatha Yoga, Shambhavi Mahamudra Kriya, Shoonya, and Samyama^3,78,79^. Shambhavi Mahamudra Kriya, a component of the Inner Engineering program, is a 21-minute practice involving sukha kriya pranayama, AUM chanting, breath fluttering, bandhas, and breath-watching meditation^3,78,81^. Sukha kriya pranayama involves gentle alternate nostril breathing, while breath watching involves observing the natural movement of breath while ignoring distractions. Shoonya meditation, meaning “no-thingness,” is an advanced practice of conscious non-doing, recommended for 15 minutes twice daily. Samyama, a 8-day retreat conducted in silence, integrates the path of awareness (Pragna) with the path of dissolution (Samadhi)^3^.

### EEG acquisition and processing

We used the 128-channel Geodesic EEG System 300 (Philips Neuro, USA) to collect data. EEG recordings took place in a dimly lit, sound-attenuated chamber with a controlled ambient temperature (25°C) and humidity (40%-60%). Electrode impedance was kept below 50 kohm as per vendor recommendations. Stimulus presentation was managed using E-prime 2.0 software (Psychology Software Tools, Inc., Sharpsburg, PA, USA). Participants were seated 90 cm from a 34 cm × 27 cm LCD monitor displaying instructions. Data were amplified with the NetAmps300 amplifier, digitized at 24-bit resolution, and sampled at 1 kHz (DC amplifier). Electrode positioning followed geodesic spacing guidelines, with Cz serving as the online reference electrode and AFz as the ground electrode.

The following steps were used in processing the EEG data:

1. Raw EEG data were imported into EEGLAB v2021.0^118^ using the Philips ’mff’ import plugin and pre-processed with in-house MATLAB scripts.
2. The data was resampled to 250 Hz and re-referenced to the average reference.
3. High-pass filtering (>0.5 Hz) and low-pass filtering (<80 Hz) were applied, along with a notch filter at 50 Hz.
4. Bad channels and segments were automatically rejected using the artifact subspace reconstruction (ASR) plugin with a 5 standard deviation threshold. On average, 11.19% (min: 1.33%, max: 45.16%) of data were removed for advanced meditators, 12.61% (min: 1.3%, max: 66.7%) for novice meditators, and 11.21% (min: 0.38%, max: 57.17%) for controls.
5. Scalp muscle, ocular, and ECG artifacts were identified and removed using the ICALABEL plugin (90% threshold) after running independent component analysis (ICA) with the ’infomax’ method. On average, 3.14 components (min: 0, max: 21) were removed for advanced meditators, 2.41 components (min: 0, max: 21) for novice meditators, and 3.99 components (min: 0, max: 25) for controls.
6. Channel interpolation was performed using neighbouring electrodes with a spherical spline approach.

### Autocorrelation window

An autocorrelation function (ACF) was calculated in MATLAB (version 2023b) using a custom code^119^. At each electrode, this function was applied to 10 second EEG epochs with 1 second sliding, across relevant session data. Within each 10 second epoch, ACF was computed on 1 second windows (with 50% overlap) and averaged to get a smooth estimate of ACF. On this, ACW-50 was computed as the first lag where the ACF decays to 50% of its maximal value^48^. Finally, the ACW-50 mean (from 20% trimmed mean) and variance (from median average deviance) values were calculated from the 10 seconds epochs for each of electrodes for each participant.

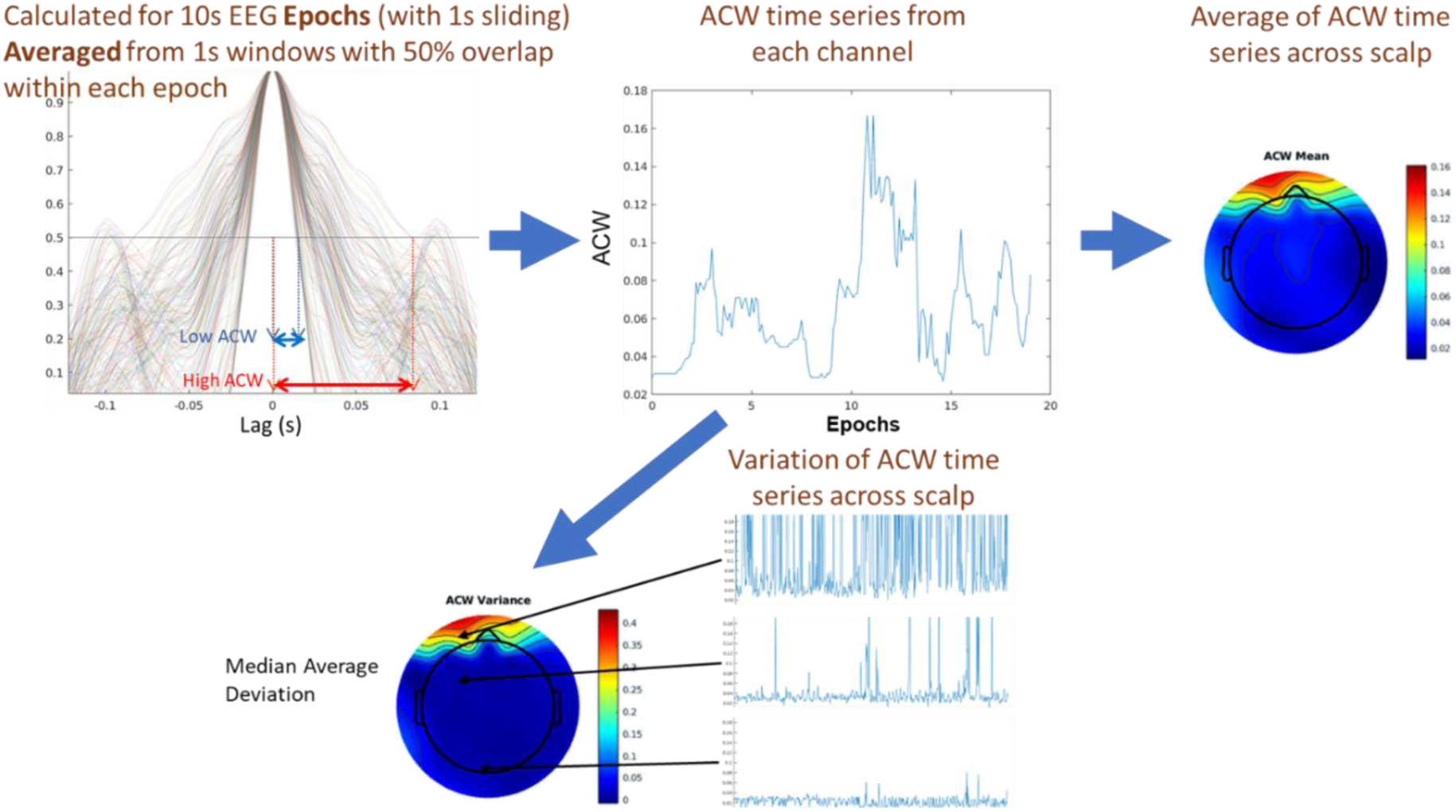
**Figure:** Steps to calculate ACW mean and variance.

### Subjective questionnaires

a. **Scale of Positive and Negative Experience (SPANE):** The SPANE (Scale of Positive and Negative Experience) consists of 12 items, divided into positive feelings (SPANE-P), negative feelings (SPANE-N), and affect balance (SPANE-B)^75^. Six items evaluate positive feelings, while the remaining six assess negative feelings. Participants rate the frequency of experiencing each item on a scale from 1 “Very rarely or never” to 5 “Very often or always.” SPANE-P and SPANE-N scores are determined by adding together the results from questions that assess positive and negative feelings, respectively. The SPANE-B score is obtained by subtracting the SPANE-N score from the SPANE-P score.
b. **Non-attachment Scale (NAS-7):** A 7-item version of the original nonattachment scale, known as the NAS-7^37^, was utilized to assess nonattachment as trait feature of the subjects. Participants rated their agreement with seven statements, such as “I can let go of regrets and feelings of dissatisfaction about the past,” using a 7-point Likert scale ranging from 1 (Strongly Disagree) to 7 (Strongly Agree).
c. **Meditation Depth Questionnaire (MEDEQ):** MEDEQ, a 30-item questionnaire, evaluates the depth of meditation practice^74^. Participants rate their level of agreement with meditation-related statements on a scale from “0 = not at all” to “4 = very much.” The total score ranges from 0 to 120, with higher scores indicating deeper meditation. The questionnaire comprises five clusters: hindrances, relaxation, concentration, essential qualities, and non-duality. MEDEQ was acquired in 70 participants (CNT – 23, NOV – 22, ADV – 25).
d. **Equanimity:** The second question of the meditation depth questionnaire, “I experienced equanimity and inner peace,” was utilized to assess equanimity. Respondents scoring 3 and 4 were categorized as having high equanimity, those scoring 2 were classified as having moderate equanimity, and those scoring 0 and 1 were categorized as having low equanimity. In addition to the single item measure of equanimity, items 2, 10, 16, and 26, as well as, even more extended, items 2, 5, 6, 10, 11, 16, 25, and 26, were utilized to assess narrow and wide equanimity scores.

The internal reliability of all the questionnaires was assessed using Cronbach’s alpha, and the calculated values were found to be between 0.8 and 0.92, indicating a high level of internal consistency.

### Data analysis

The statistical analyses were conducted using RStudio Version 1.4.1106, employing several R packages including ggplot2^120^, ggsci^121^, ggstatsplot^122^, dplyr^123^, effectsize^124^, performance^125^, and car^126^ for necessary analyses. Non-parametric tests such as Kruskal-Wallis, pairwise Dunn test, Mann-Whitney, and chi-squared test were utilized as the data did not meet parametric test assumptions. Effect sizes were reported with 95% confidence intervals, and significance was set at p < 0.05, with exact p-values provided. Statistical reporting followed the guidelines of the American Psychological Association^127^, as implemented in the ggstatsplot package^122^. Partial mediation analysis was run in R using lavaan package^128^. p value was estimated via bootstrap statistics (n=5000). 95% CI was also estimated. The mediation effect was captured as a product of indirect paths. In the EEG data, we carried out Mann-Whitney tests to compare the ACW mean and variance values between controls and advanced meditators during breath watch and task. In situations where there was no significance found (like between breath watch and task in advanced meditators), we tested for non-inferiority. Spearman’s rho method was used to estimate the correlation value as the questionnaires were ordinal in nature. Correlation analysis was done in R using the cor.test function with exact=FALSE to compute p values when there are ties between the ranks. Visualisation was done using the ggplot library and each dot colored as per group. A line of fit was drawn with 95% CI shadow.

## Notes

### Competing Interest Statement

The authors have declared no competing interest.

