## Supplementary file for "The Balanced Mind and its Intrinsic Neural Timescales in Advanced Meditators"

### Supplementary information

**Table 1 Criteria to classify study participants into advanced, novice, and controls**

| Participants | Criteria |
|---|---|
| **Advanced meditators** | • Must have completed advanced 8-day meditation retreat (Samyama).<br>• No history or current practice of any other form of Yoga or meditation<br>• Must be practicing various Isha Yoga practices such as hatha yoga, Shambhavi Mahamudra, Shakti Chalana Kriya, Shoonya, and Samyama meditation daily. |
| **Novice meditators** | • Must have been initiated into Shambhavi Mahamudra Kriya and regularly practicing.<br>• No history or current practice of any other form of Yoga or meditation.<br>• Must be practicing Isha Yoga daily. |
| **Meditation-naïve controls** | • No prior exposure to any meditation or yoga |

**Table 2 Inclusion and Exclusion criteria**

| Inclusion criteria | Exclusion criteria |
|---|---|
| a) Male and Female participants | a) History of neurological disease, vision defects (uncorrected), auditory deficits or serious physical disabilities |
| b) Age-range 25-50 years | b) History of substance abuse/dependence |
| c) Healthy or under stable dosage of non-psychoactive medication (stable dosage period considered as a minimum of one month with no changes anticipated). | c) History of major mental illness, any psychiatric medication, psychotherapy |
| d) Able to read, write, and speak fluently in English. Able to comprehend and answer standard questionnaires. | |
| e) Able and willing to cooperate in EEG studies | |

f) Informed Consent as approved by

Institute Human Ethics Committee

**Instructions to participants**

**Rest**

Instructions were given to the controls and meditators to sit still and relax. Prior instructions were given to avoid unnecessary movements as high-frequency brain oscillations, especially in the gamma band, are very sensitive to muscular movements, and interfere with analysis.

**Pranayama**

Isha meditators and controls performed a form of alternate nostril breathing pranayama called Sukha Kriya pranayama and Nadi Shuddhi, respectively.

**Breath Watching**

Isha meditators practiced breath-watching as per tradition.

Following instructions were given to controls: Pay attention to your breath. When the mind wanders, simply notice that the mind has wandered and bring attention back to the breath.

**Shoonya meditation**

The following instructions were given to controls:

Sit still. Do not do anything in particular. Ignore mental activity. Neither suppress any thought nor follow any thought. Isha meditators practiced Shoonya as they were initiated into it.

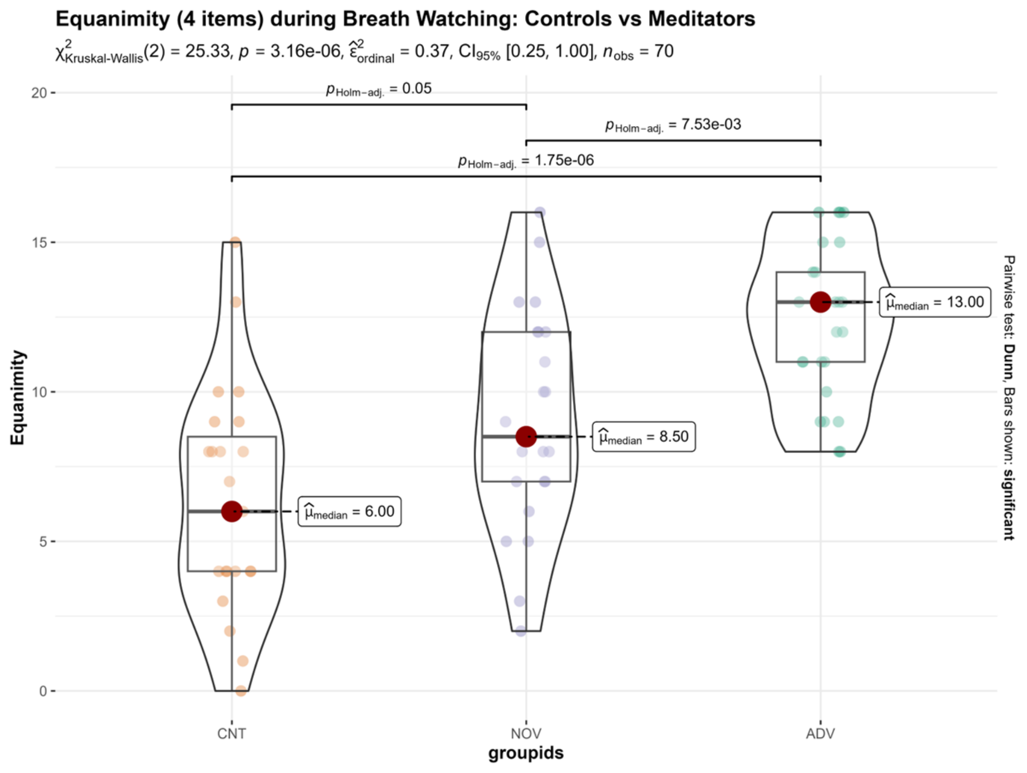


**Supplement Figure 1:** Equanimity (4 items) during breath watch.

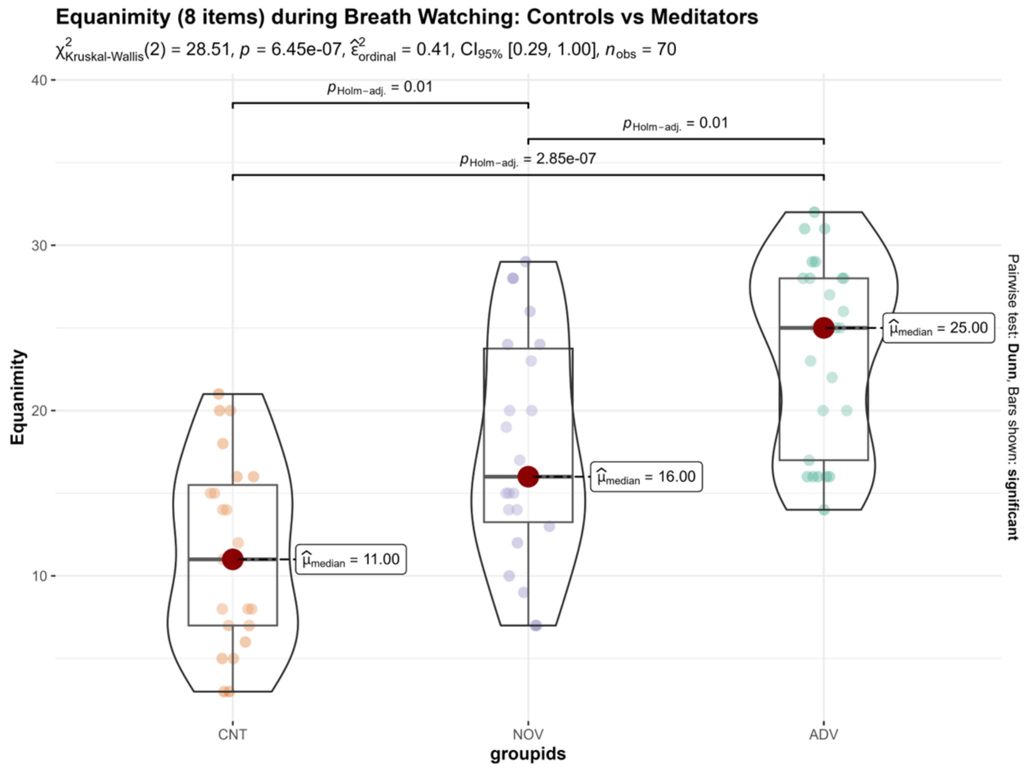


**Supplement Figure 2:** Equanimity (8 items) during breath watch.

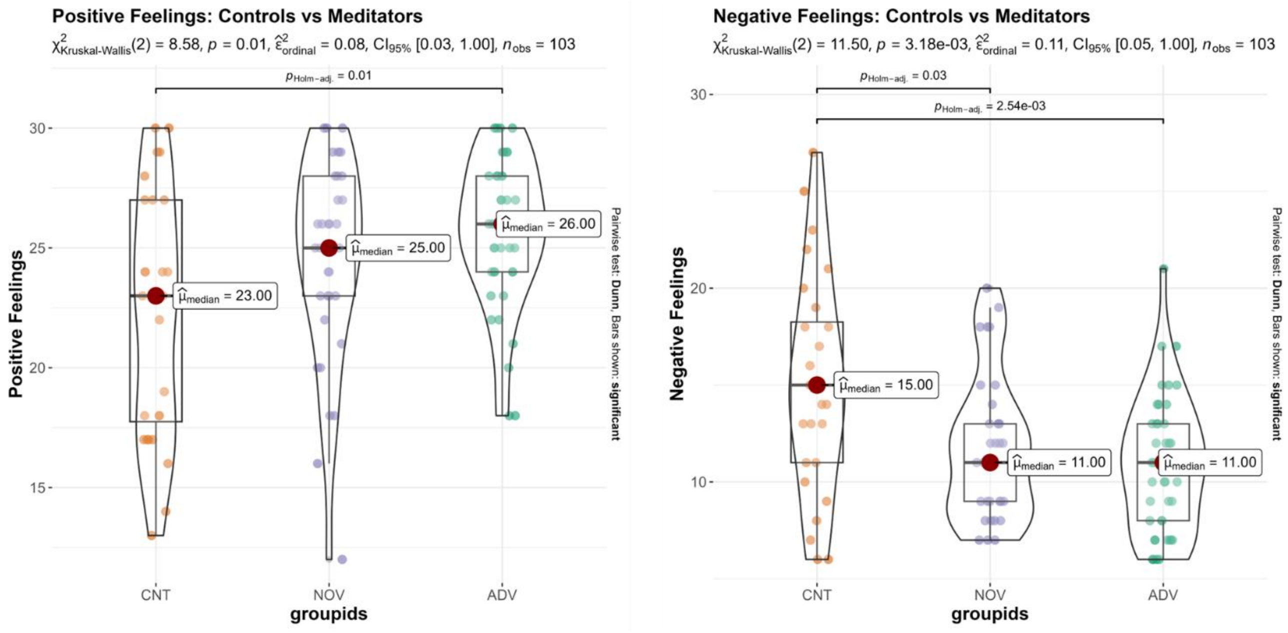


**Supplement Figure 3:** Positive and negative feelings in controls and meditators.

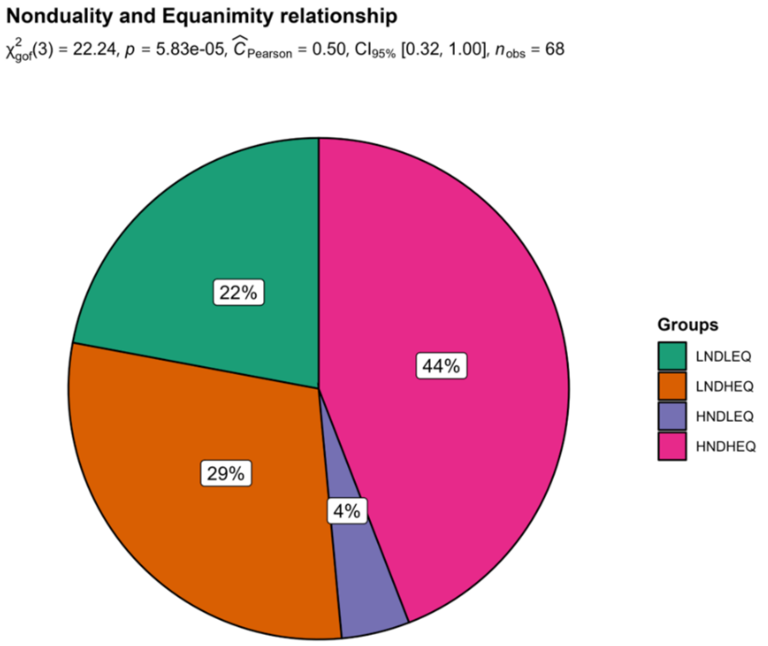


**Supplement Figure 4:** Relationship between nonduality and equanimity. LNDLEQ: Low Non-duality Low Equanimity, LNDHEQ: Low Non-duality High Equanimity, HNDLEQ: High Non-duality Low Equanimity, and HNDHEQ: High Non-duality High Equanimity.

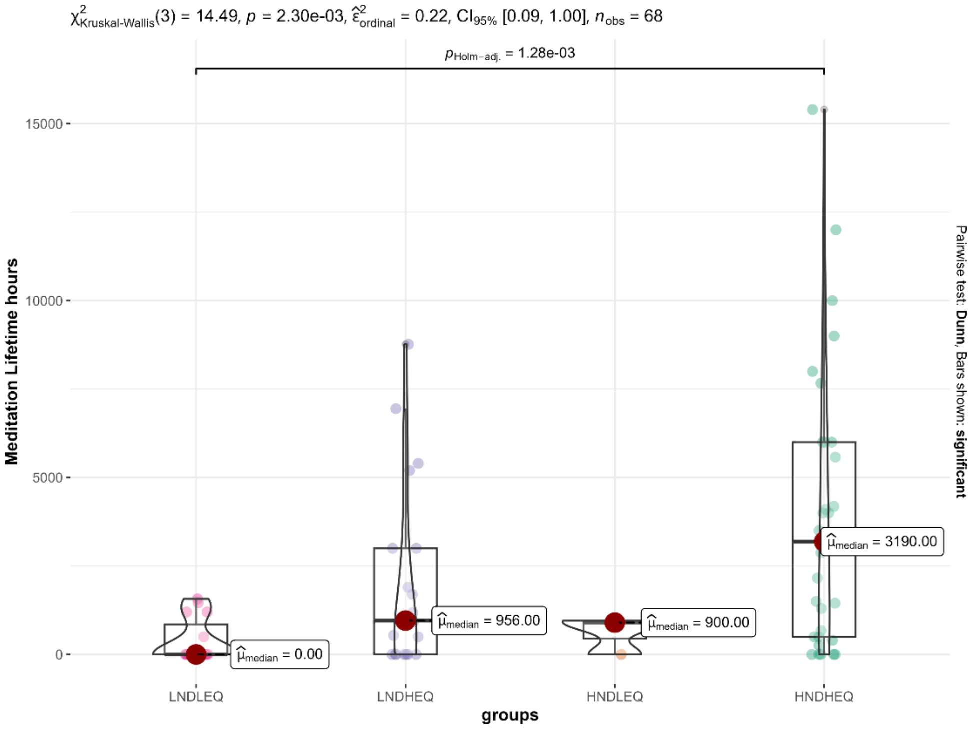


**Supplement Figure 5:** Relationship between four groups and meditation lifetime hours. LNDLEQ: Low Non-duality Low Equanimity, LNDHEQ: Low Non-duality High Equanimity, HNDLEQ: High Non-duality Low Equanimity, and HNDHEQ: High Non-duality High Equanimity.

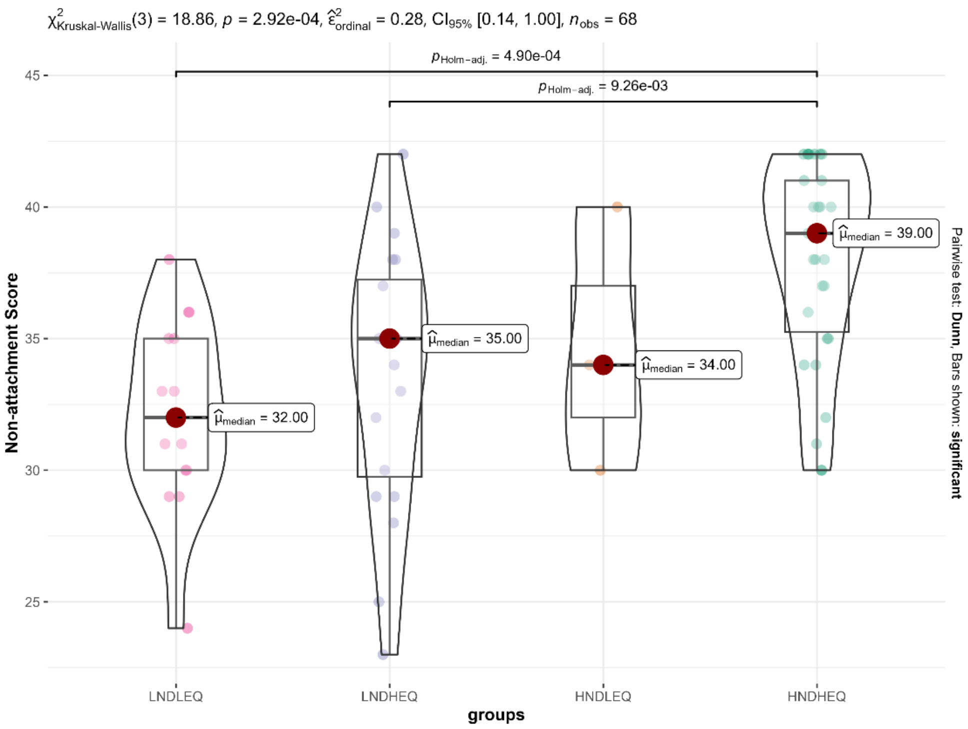

**Supplement Figure 6:** Relationship between four groups and non-attachment. LNDLEQ: Low Non-duality Low Equanimity, LNDHEQ: Low Non-duality High Equanimity, HNDLEQ: High Non-duality Low Equanimity, and HNDHEQ: High Non-duality High Equanimity.

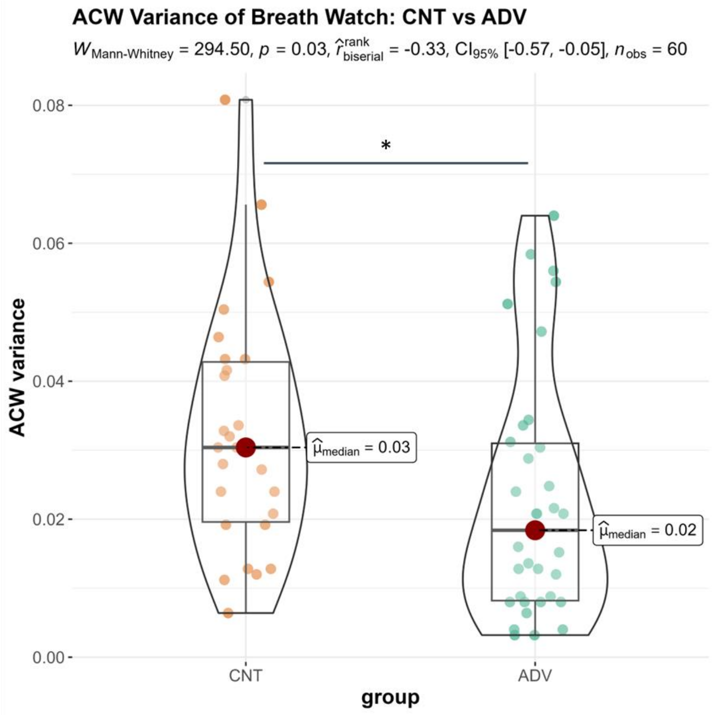


**Supplement Figure 7:** ACW variance during breath watch between controls and advanced meditators.

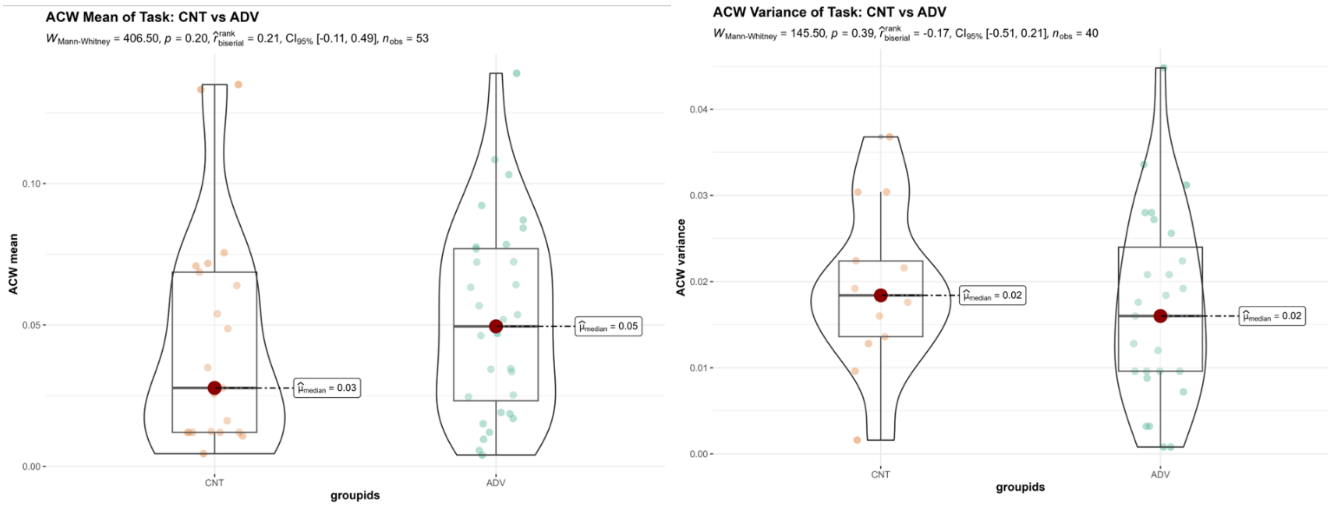


**Supplement Figure 8:** ACW mean (left) and variance (right) during task between controls and advanced meditators.

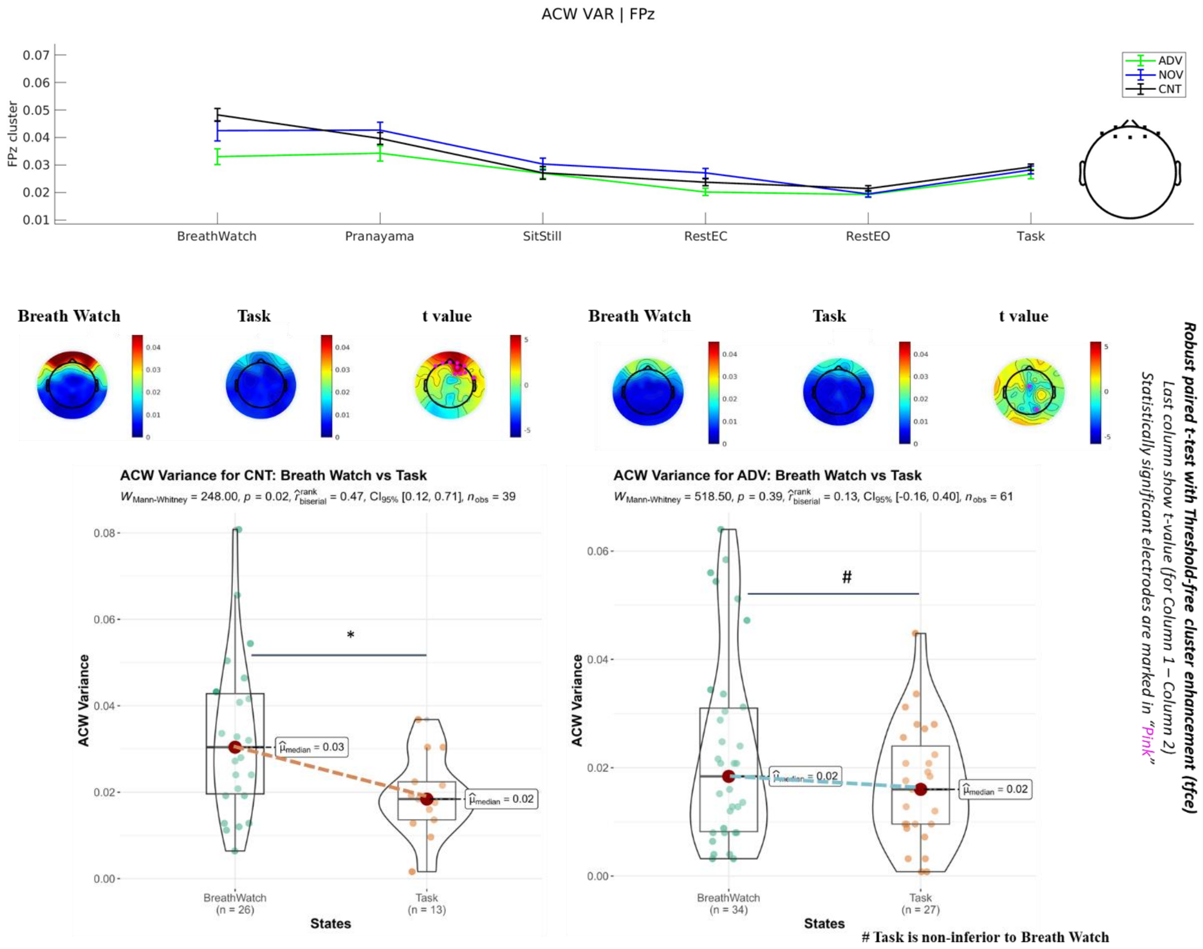


**Supplement Figure 9:** The top panel shows the line plot for the ACW variance (FpZ cluster, top right) along the internal-external attention gradient. The middle panel shows the topoplots of the ACW variance during breath watch and task for controls (left) and advanced meditators (right). The last column of the topoplots shows t-values (column 1 – column 2) and statistically significant electrodes are marked in pink. The bottom panel shows the ACW variance for controls (left) and advanced meditators (right) for breath watch vs task.

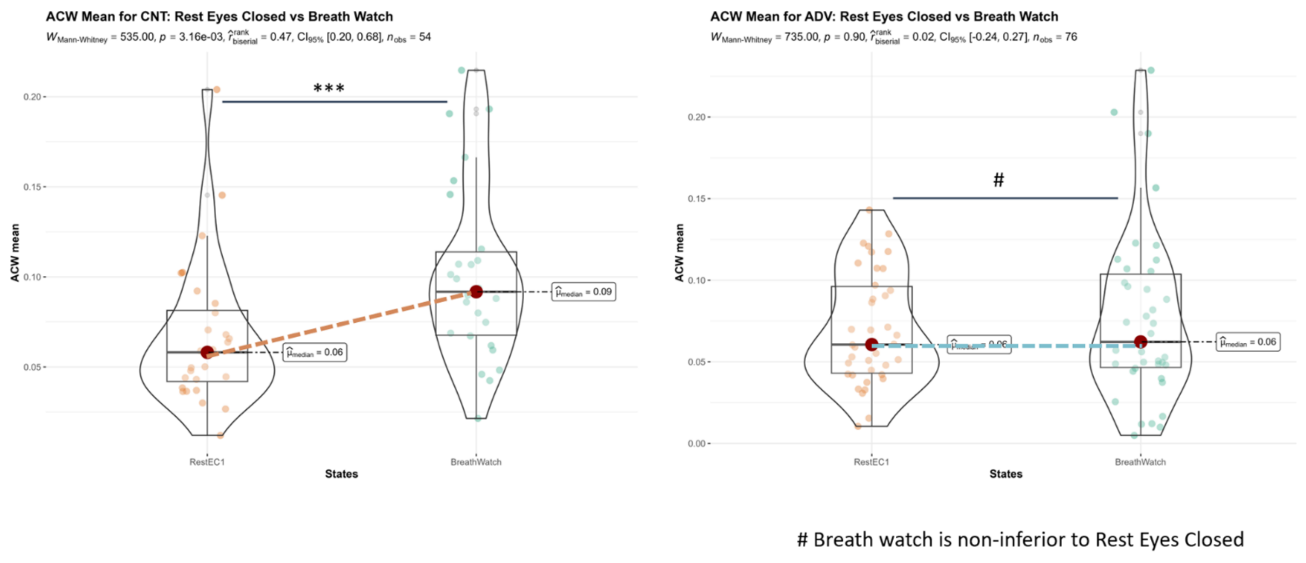


**Supplement Figure 10:** ACW mean for rest eyes closed vs breath watch for controls (left) and advanced meditators (right)

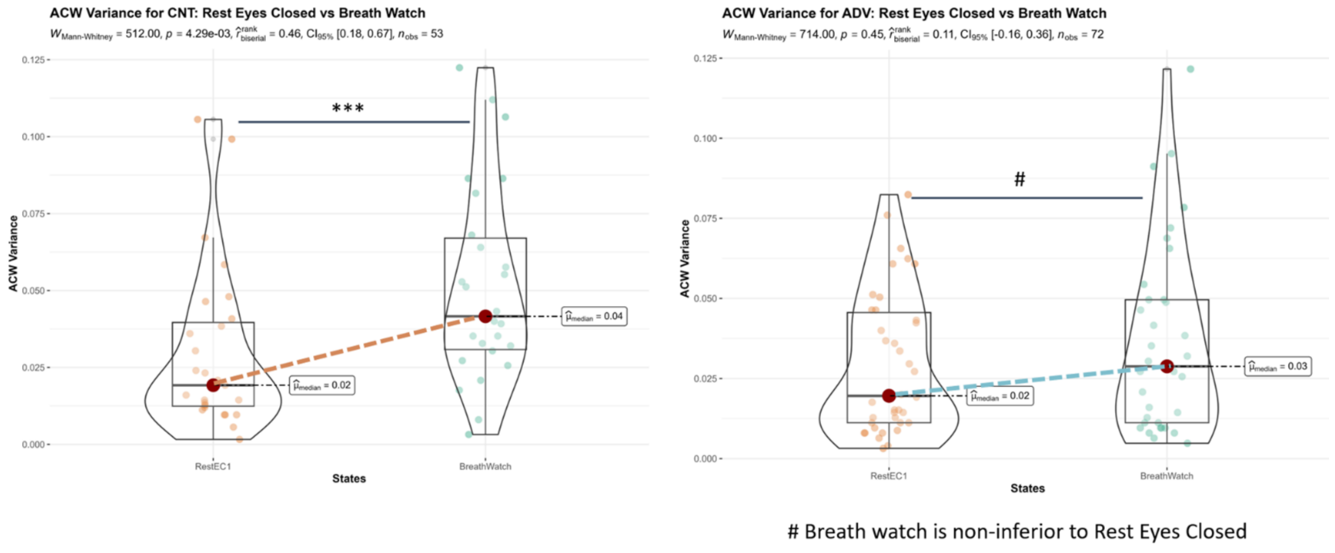


**Supplement Figure 11:** ACW variance for rest eyes closed vs breath watch for controls (left) and advanced meditators (right)

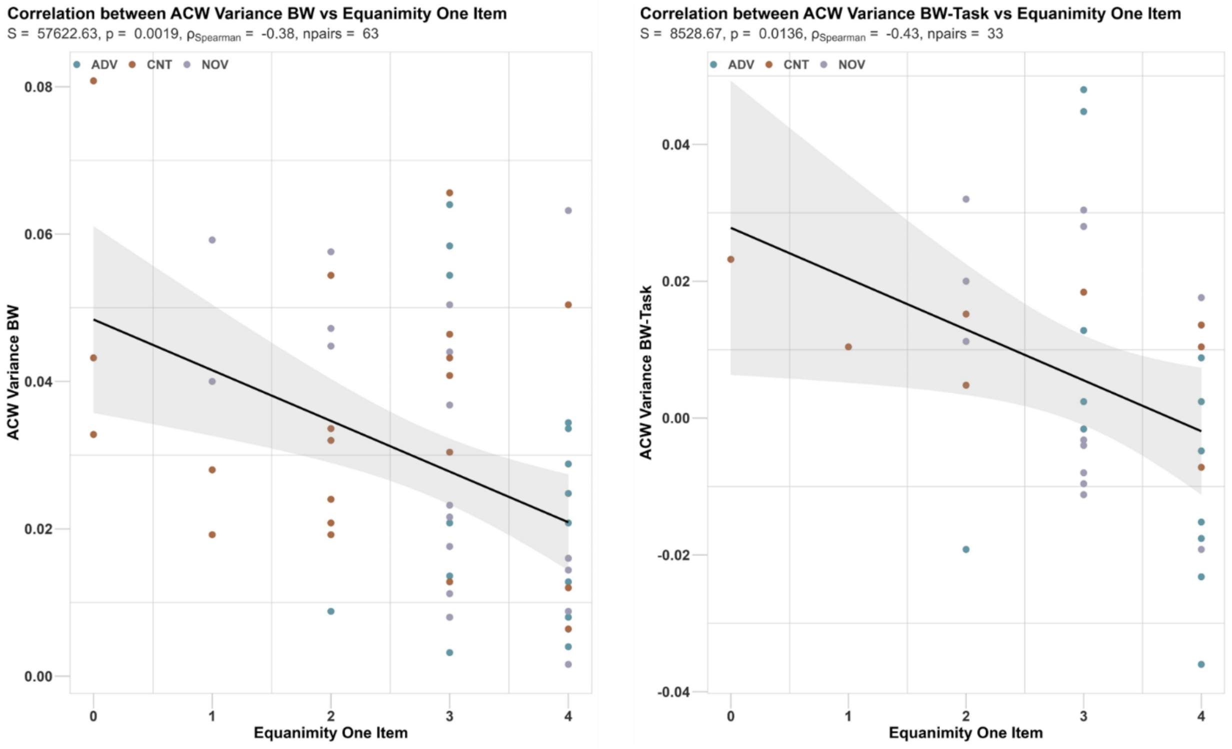


**Supplement Figure 12:** Correlations between ACW variance and Equanimity. Shadows show 95% CI intervals.

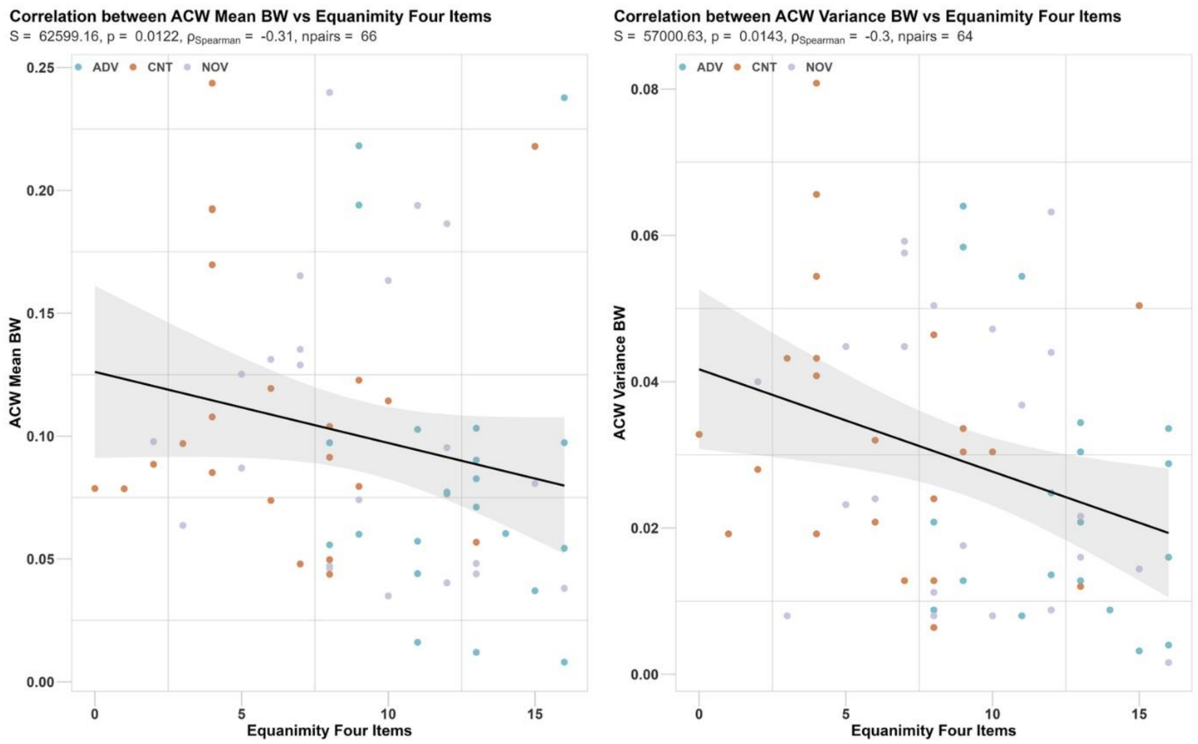


**Supplement Figure 13:** Correlation between meditation Equanimity four items and ACW mean (left) and ACW variance (right)

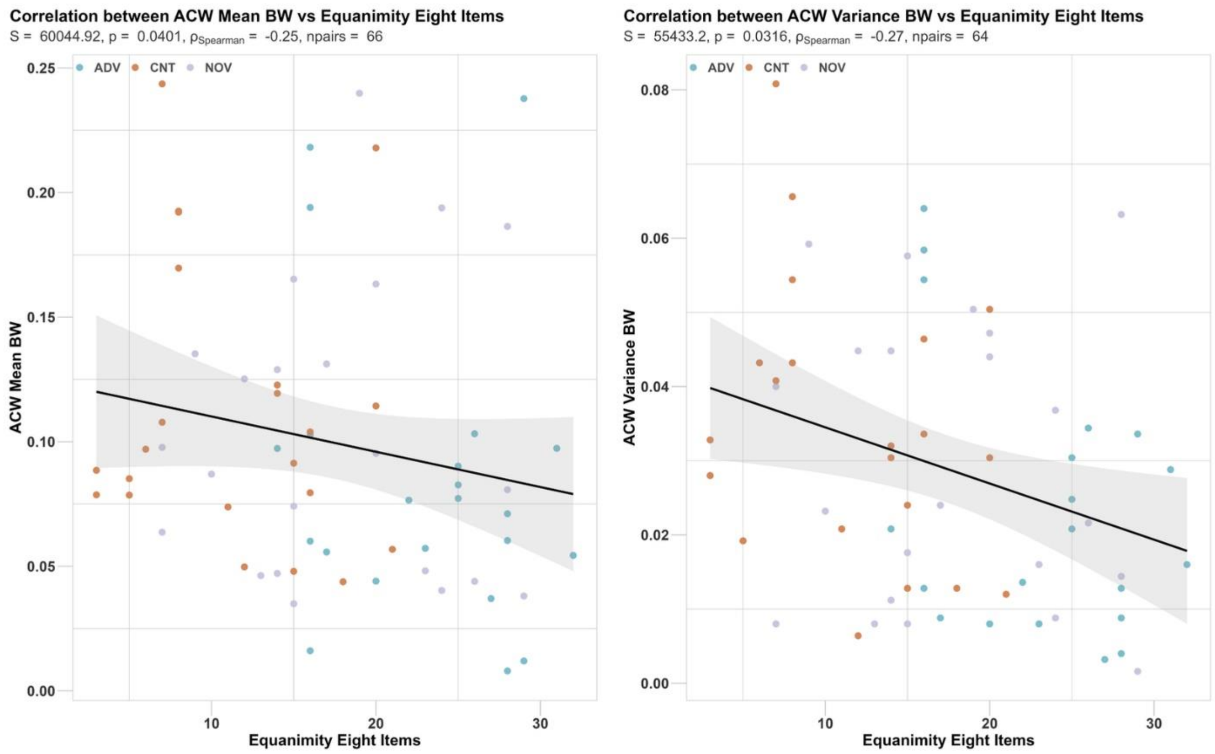


**Supplement Figure 14:** Correlation between meditation Equanimity eight items and ACW mean (left) and ACW variance (right)
